# Solving the Sample Size Problem for Resource Selection Analysis

**DOI:** 10.1101/2021.02.22.432319

**Authors:** Garrett M. Street, Jonathan R. Potts, Luca Börger, James C. Beasley, Stephen Demarais, John M. Fryxell, Philip D. McLoughlin, Kevin L. Monteith, Christina M. Prokopenko, Miltinho C. Ribeiro, Arthur R. Rodgers, Bronson K. Strickland, Floris M. van Beest, David A. Bernasconi, Larissa T. Beumer, Guha Dharmarajan, Samantha P. Dwinnell, David A. Keiter, Alexine Keuroghlian, Levi J. Newediuk, Júlia Emi F. Oshima, Olin Rhodes, Peter E. Schlichting, Niels M. Schmidt, Eric Vander Wal

**Affiliations:** Department of Wildlife, Fisheries, and Aquaculture, Mississippi State University, Mississippi State, MS, 39762, USA; Quantitative Ecology and Spatial Technologies Laboratory, Mississippi State University, Mississippi State, MS, 39762, USA; School of Mathematics and Statistics, University of Sheffield, Sheffield, S3 7RH, United Kingdom; Department of Biosciences, Swansea University, Swansea, SA2 8PP, United Kingdom; Centre for Biomathematics, Swansea University, Swansea, SA2 8PP, United Kingdom; Savannah River Ecology Laboratory, University of Georgia, Aiken, SC, 29802, USA; Department of Integrative Biology, University of Guelph, Guelph, Ontario, N1G 2W1, Canada; Department of Biology, University of Saskatchewan, Saskatoon, Saskatchewan, S7N 5E2, Canada; Haub School of Environment and Natural Resources, University of Wyoming, Laramie, WY, 82072, USA; Department of Biology, Memorial University of Newfoundland, St. John’s, Newfoundland, A1B 3X9, Canada; Instituto de Biosciências, Universidad Estadual Paulista, Rio Claro, São Paulo, Brazil; Centre for Northern Forest Ecosystem Research, Ontario Ministry of Natural Resources and Forestry, Thunder Bay, Ontario, P7E 6S8, Canada; Department of Bioscience, Aarhus University, Aarhus, Denmark; Wyoming Cooperative Fish and Wildlife Research Unit, University of Wyoming, Laramie, WY, 82071, USA; IUCN/SSC Peccary Specialist Group, Campo Grande, Brazil

**Keywords:** bootstrap, habitat selection, p-value, power analysis, Resource Selection Function, sample size, Species Distribution Model, validation

## Abstract

1. Sample size sufficiency is a critical consideration for conducting Resource-Selection Analyses (RSAs) from GPS-based animal telemetry. Cited thresholds for sufficiency include a number of captured animals *M* ≥ 30 and as many relocations per animal *N* as possible. These thresholds render many RSA-based studies misleading if large sample sizes were truly insufficient, or unpublishable if small sample sizes were sufficient but failed to meet reviewer expectations.
2. We provide the first comprehensive solution for RSA sample size by deriving closed-form mathematical expressions for the number of animals *M* and the number of relocations per animal *N* required for model outputs to a given degree of precision. The sample sizes needed depend on just 2 biologically meaningful quantities: habitat selection strength and a novel measure of landscape complexity, which we define rigorously. The mathematical expressions are calculable for any environmental dataset at any spatial scale and are applicable to any study involving resource selection (including sessile organisms). We validate our analytical solutions using globally relevant empirical data including 5,678,623 GPS locations from 511 animals from 10 species (omnivores, carnivores, and herbivores living in boreal, temperate, and tropical forests, montane woodlands, swamps, and arctic tundra).
3. Our analytic expressions show that the required *M* and *N* must decline with increasing selection strength and increasing landscape complexity, and this decline is insensitive to the definition of availability used in the analysis. Our results contradict conventional wisdom by demonstrating that the most biologically relevant effects on the utilization distribution (i.e. those landscape conditions with the greatest absolute magnitude of resource selection) can often be estimated with far fewer data than is commonly assumed.
4. We identify several critical steps in implementing these equations, including (i) a priori selection of expected model coefficients, and (ii) sampling intensity for background (absence/pseudo-absence) data within a given definition of availability. We show that random sampling of background data violates the underlying mathematics of RSA, leading to incorrect values for necessary *M* and *N* and potentially incorrect RSA model outputs. We argue that these equations should be a mandatory component for all future RSA studies.

## Introduction

Resource selection analysis (RSA) is a broad framework linking the distribution of animals to their preferences for specific habitat conditions and is a fundamental tool in animal ecology (Boyce & McDonald 1999; Strickland & McDonald 2006). Obtaining sufficient locations to ascertain the distribution of animals across landscapes is a fundamental requirement for RSA. Indeed, to understand intra-specific variation in the distribution of animals – a critical research aim in basic and applied animal ecology – it is necessary to obtain repeated localizations on multiple individuals, now commonly collected using animal-attached GPS sensors (Hebblewhite & Haydon 2010). GPS data on animal movements are hence commonly employed for RSA and are often analyzed using Resource Selection Functions (RSFs; Boyce & McDonald 1999; Manly *et al.* 2002; Elith & Leathwick 2009; Hebblewhite & Haydon 2010). RSFs are a class of exponential models of space use that estimate the probability distribution of animal locations using different resources/conditions in the landscape, taking into account the availability of each resource, and thereby provide a measure of the ‘strength’ of (behavioral) selection for or against each resource (Manly *et al.* 2002). RSFs are easily fitted using standard statistical models (commonly logistic or conditional logistic regression) applied to data on animal locations and resource distributions in the landscape and have become a cornerstone of research in spatial ecology (Manly *et al.* 2002; Elith & Leathwick 2009; Renner & Warton 2013).

Given the prevalence of RSFs, it is surprising that the central question determining the validity of inferences obtained – how much data is needed to estimate a RSF for a given species? – has not been solved. This issue has been broached for occupancy analysis (Guillera-Arroita & Lahoz-Monfort 2012) and generalized linear mixed models (Johnson *et al.* 2015), and has been evaluated within individual RSF studies using simulations (Leban *et al.* 2001; Loe *et al.* 2012), yet no analytic expressions exist to determine the number of animals (*M*) and relocations per animal (*N*) required to obtain RSF outputs to a given degree of precision. While the accuracy and precision of RSFs generally increase with sample size, leading to a standard rule-of-thumb of *M* ≥ 30 needed for reliable ecological inference (Leban *et al.* 2001), this rough guideline is grounded in century-old thinking about statistics in the pre-computation world (James *et al.* 2013). Crucially, it is also oblivious to the ecological reality that a multitude of factors may affect selection strength and determine the required sample size (Manly *et al.* 2002; McLoughlin *et al.* 2010; Hebblewhite & Haydon 2010). These include density-dependence (i.e. certain habitats become less attractive when occupied by conspecifics; Fretwell & Lucas 1969; McLoughlin *et al.* 2010; van Beest *et al.* 2016), trade-offs in selection for forage and cover under predation risk (Fortin *et al.* 2005; McLoughlin *et al.* 2010), temporal variations in resource dynamics (McLoughlin *et al.* 2010; Paolini *et al.* 2018), or the degree of habitat availability or heterogeneity in a landscape (Mysterud & Ims 1998; McLoughlin *et al.* 2010; van Beest *et al.* 2016; Paolini *et al.* 2018). There is no consistency in RSF studies in the number of replicates used (Hebblewhite & Haydon 2010), as the only alternative approaches to establishing the number of replicates a priori are ecologically informed guesswork, or simply to collect as much data as possible.

The crux of the problem lies in the relationship between sample size and ecological complexity. It is suggested that more complex systems require more data to describe (Wisz *et al.* 2008), yet a robust power analysis (Johnson *et al.* 2015) allowing examination of the relationship between RSF estimation, system complexity, and data availability is crucially missing. This has obvious economic and ethical implications if more animals are tagged and monitored than needed and affects research aimed at the conservation of species, which requires reliable estimates of animal-habitat relationships but where it is often impossible to monitor large numbers of animals. Here, we provide a solution to the sample size problem in RSFs by deriving analytic expressions for the values of *M* and *N* (the number of animals and relocations per animal respectively) required to estimate RSFs to a required degree of accuracy, taking into account landscape complexity and the strength of selection for the resources. We validate these expressions using simulations and a large dataset of GPS-tagged animals (including 10 species from different continents and biomes) and show that the most biologically relevant effects of landscapes on animal distributions can often be estimated with far fewer animals and locations than are commonly stated.

## Methods

We begin by describing mathematically how to determine the number of locations per animal (*N*) and the number of animals (*M*) for RSA. RSA seeks to parametrize a model of space use that has the following form (Manly *et al.* 2002):

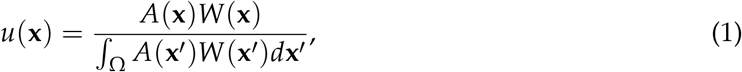

where *u*(**x**) is the *utilization distribution* of the study species (i.e. the probability density function of the study animals’ locations), *A*(**x**) is a function denoting the availability of the point **x** to the animals, Ω is the study area, and *W*(**x**) is the RSF. (Note: throughout this manuscript, bold fonts imply that the quantity is a vector.) For the purposes of our analytic calculations, our RSF will be dependent upon a single resource layer *R*(**x**). This could denote, for example, the vegetation quality or prey availability at point **x**. However, in general, *R*(**x**) represents a map of any environmental feature which is hypothesized to covary with space use. Although we only look at one resource layer at a time for our analytic calculations, we show in our empirical study (below) that the resulting formulae work when the RSF has multiple layers.

As is the standard method for RSA, we make 3 simplifying assumptions (Manly *et al.* 2002): (i) our weighting function is of the form *W*(**x**|*β*) = exp[*βR*(**x**)], where *β* is a parameter to be estimated; (ii) the availability kernel *A*(**x**) is a uniform distribution; and (iii) relocations are independent. Consequently, our model of space use from Equation (1) becomes:

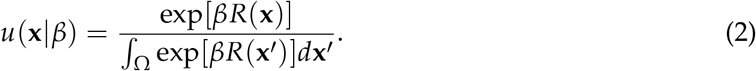

The aim of this section is to understand how many independent samples are required to give an accurate parametrization of the model in Equation (2).

### Locations from a Single Individual (*N*)

We first need to phrase the question “How many locations?” in a concrete, mathematical way. Suppose we wish to test the null hypothesis *H*_0_ : *β* = 0 against the alternative *H*_1_ : *β* ≠ 0 at a significance level *p* ∈ (0, 1). An experiment to test this hypothesis involves measuring *N* samples and using (conditional) logistic regression to infer *β* and test the null hypothesis (as is the standard method for resource selection, e.g. Manly *et al.* 2002). We define *N*_*α,p*_(*β*) to be the minimum number of samples required so that we expect to reject the null hypothesis in 100(1 − *α*)% of experiments. An approximate analytical formula for *N*_*α,p*_(*β*) is given as follows (derived in Supplementary Appendix A):

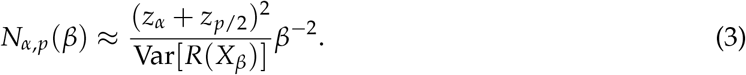

Here, *z*_*α*_ = Φ^−1^(1 − *α*) where Φ(·) is the cumulative distribution function for the standard normal distribution (e.g. *z*_0.05_ ≈ 1.645, *z*_0.025_ ≈ 1.96), *X*_*β*_ is a random variable whose probability density function is given by Equation (2), and Var[*R*(*X*_*β*_)] is the variance of *R*(*X*_*β*_). An explicit functional expression for Var[*R*(*X*_*β*_)] can be written as follows:

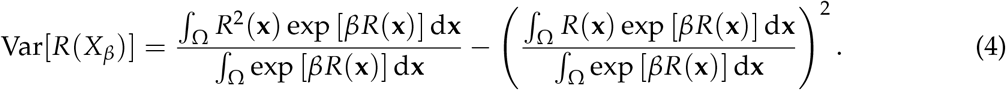

We call Var[*R*(*X*_*β*_)] “landscape complexity”. Critically, this form of landscape complexity is determined in part by multiplying the landscape layer by the expected *β*, so it should be understood as representing the landscape complexity *as viewed by the animal*.

The formula in Equation (3) is approximate due to two assumptions: (i) it relies on the standard error, *σ*, of the maximum likelihood function being approximately normally distributed, and (ii) it uses a standard result relating the standard error for the estimator of *β* to the second derivative of the log-likelihood function (see Supplementary Appendix A for more details). Therefore it is necessary to investigate the magnitude of these approximating assumptions using simulated data.

To test how effective the approximate expression from Equation (3) is at capturing the actual number of samples required to infer *β* with a given level of accuracy, we constructed a simulated resource layer which describes an example of the function *R*(**x**) (Fig. 1a). This test layer is a Gaussian random field, previously used in the context of resource selection by Potts *et al.* (2014). It was generated by the R function `GaussRF()` from the `RandomFields` package (Schlather *et al.*, 2016), using the exponential model with mean=0, variance=1, nugget=0, and scale=10, and consists of *L* = 100 by *L* = 100 pixels. By sampling *N* times from Equation (2) for various *N* with *R*(**x**), we can compute empirical values for *N*_*α,p*_(*β*) for different *β* (full method given in Supplementary Appendix B). Comparison of these empirically-derived values alongside the analytical expression from Equation (3) reveals remarkably strong agreement (Fig. 1b). This suggests that Equation (3) gives an accurate estimation of the number of independent samples required to estimate *β*.

**Figure 1:**
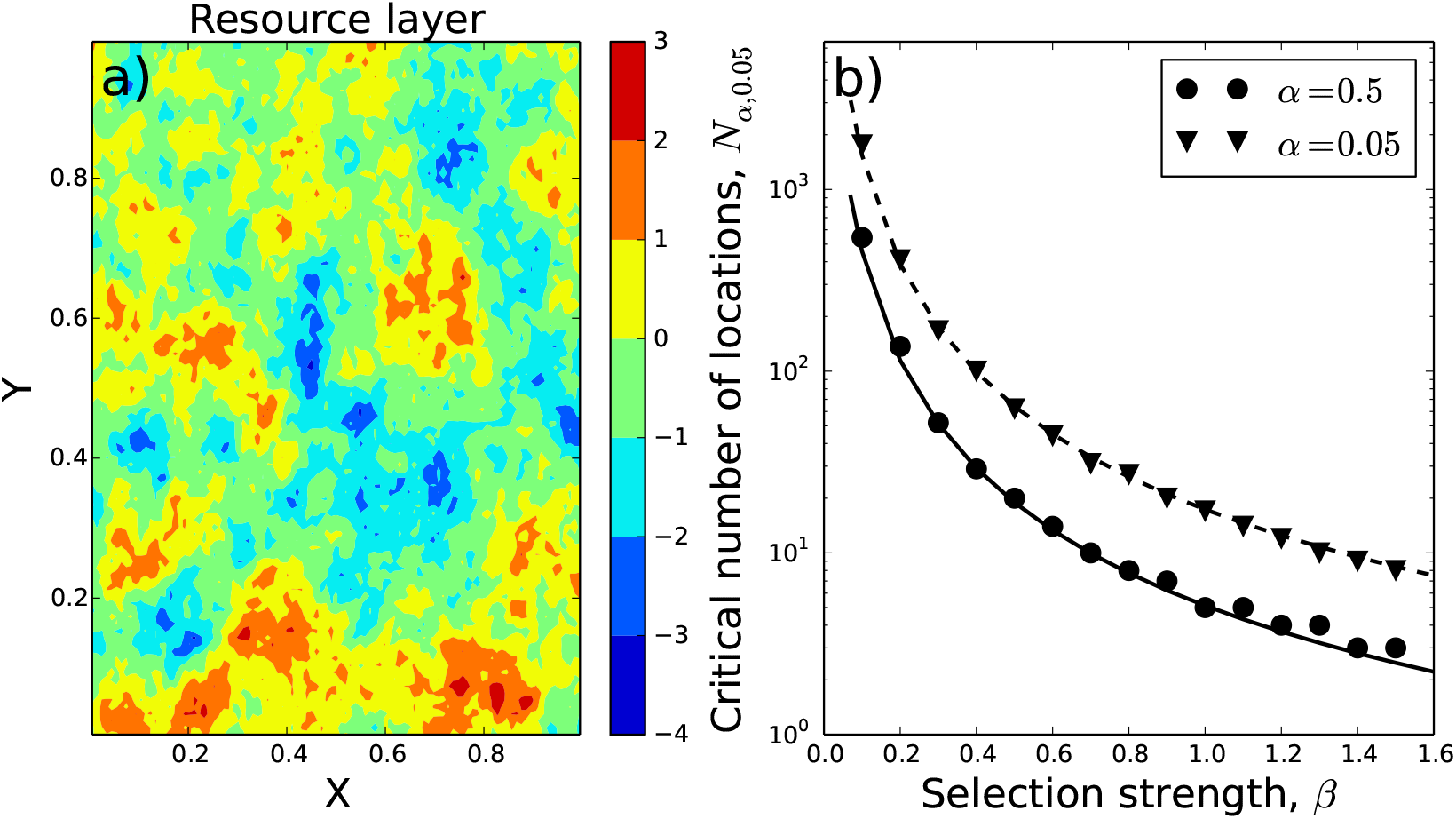
Performance of analytic expression on simulated data. Panel (a) shows a simulated resource layer, *R*(**x**), which was used to construct the utilisation distribution (Equation 2) from which the simulated animal locations were samples. The circles (resp. triangles) in Panel (b) show the empirically-derived values of *N*_0.5,0.05_(*β*) (resp. *N*_0.05,0.05_(*β*)), the minimum number of samples required so that there is a 50% chance (resp. 95% chance) of rejecting the null hypothesis that *β* = 0 at a significance level of *p* = 0.05. The solid line (resp. dashed line) in Panel (b) shows the corresponding analytic approximations given by Equation (3) and the remarkable agreement with the empirically-derived values.

### Locations from multiple individuals (*M*)

Now we assume that there are *M* individuals and they each select resources with different *β*. To model this, let *β*_1_, ..., *β*_*M*_ ~ *N*(*β*, *s*^2^) be independent draws from a normal distribution with mean *β* and variance *s*^2^. Then *β*_*i*_ is the coefficient of resource selection for individual *i* ∈ {1, ..., *M*}. Suppose for each individual *i* we have gathered *N*_*i*_ locations. Let 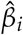 be the maximum likelihood estimator for *β*_*i*_. Then the standard deviation of 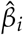 can be estimated as (Supplementary Appendix A, Equation 15):

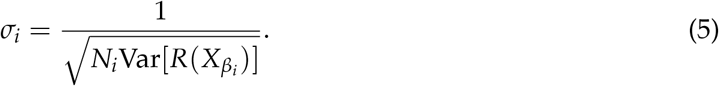

If 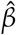 is the mean of 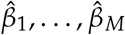, then 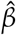 is normally distributed as follows (Supplementary Appendix C):

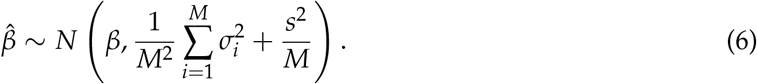

Thus 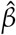 is an unbiased estimator of *β*. Notice that the variance decays as *M* increases. If the practitioner has some prior expectation of the possible values of *β* and *s*^2^, Equation (6) can be used to calculate the number of animals, *M*, required to obtain an empirical estimate of *β* to a given degree of accuracy.

As well as calculating an estimate of *β*, it is also possible to estimate *s*^2^. The following is an unbiased estimator of *s*^2^ for *M* ≥ 2 (Supplementary Appendix C):

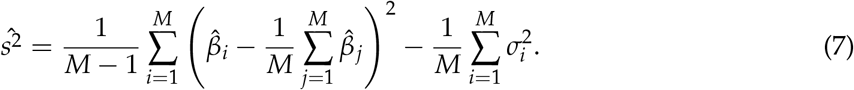

We were not able to derive a closed analytic formula for the uncertainty in the estimator given in Equation (7); however, we provide code for estimating this using random sampling (see Supplementary Appendix D). In general, the estimator becomes more precise for lower *σ*_*i*_ and higher *M*. This is shown in Supplementary Appendix D, where we also verify numerically Equations (6) and (7).

Equation (6) allows us to calculate the minimum number of animals, *M*_*α,p*_(*β*), for which we would expect to reject the null hypothesis that *β* = 0, at significance level *p*, 100(1 − *α*)% of the time (two-tailed test). *M*_*α,p*_(*β*) is the minimum integer, *M*, that satisfies the following inequality:

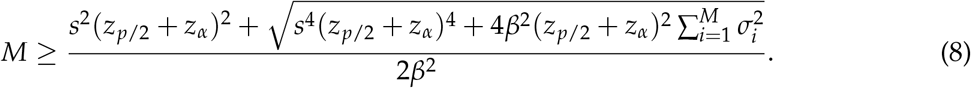

### Data and Resource Selection Functions

Equations (3) and (8) give predicted values for the number of relocations *N* and the number of animals *M* required for RSF estimation. To test our analytical predictions, we compiled GPS-based relocation datasets from 10 separate species with accompanying landscape data in raster format (Table S1; Fig. 2). Landscape data were either categorical (i.e. discrete landscover) or numeric (e.g. elevation, precipitation, etc.). To ensure comparability between model outputs for each species, we centered and scaled each numeric landscape raster in R using the `scale()` function with default parameters. We converted categorical landcover rasters to binary raster layers for each landcover classification of interest (e.g. deciduous forest, croplands, etc.) to acquire estimates of Var[*R*(*X*_*β*_)] for a given categorical raster.

**Figure 2:**
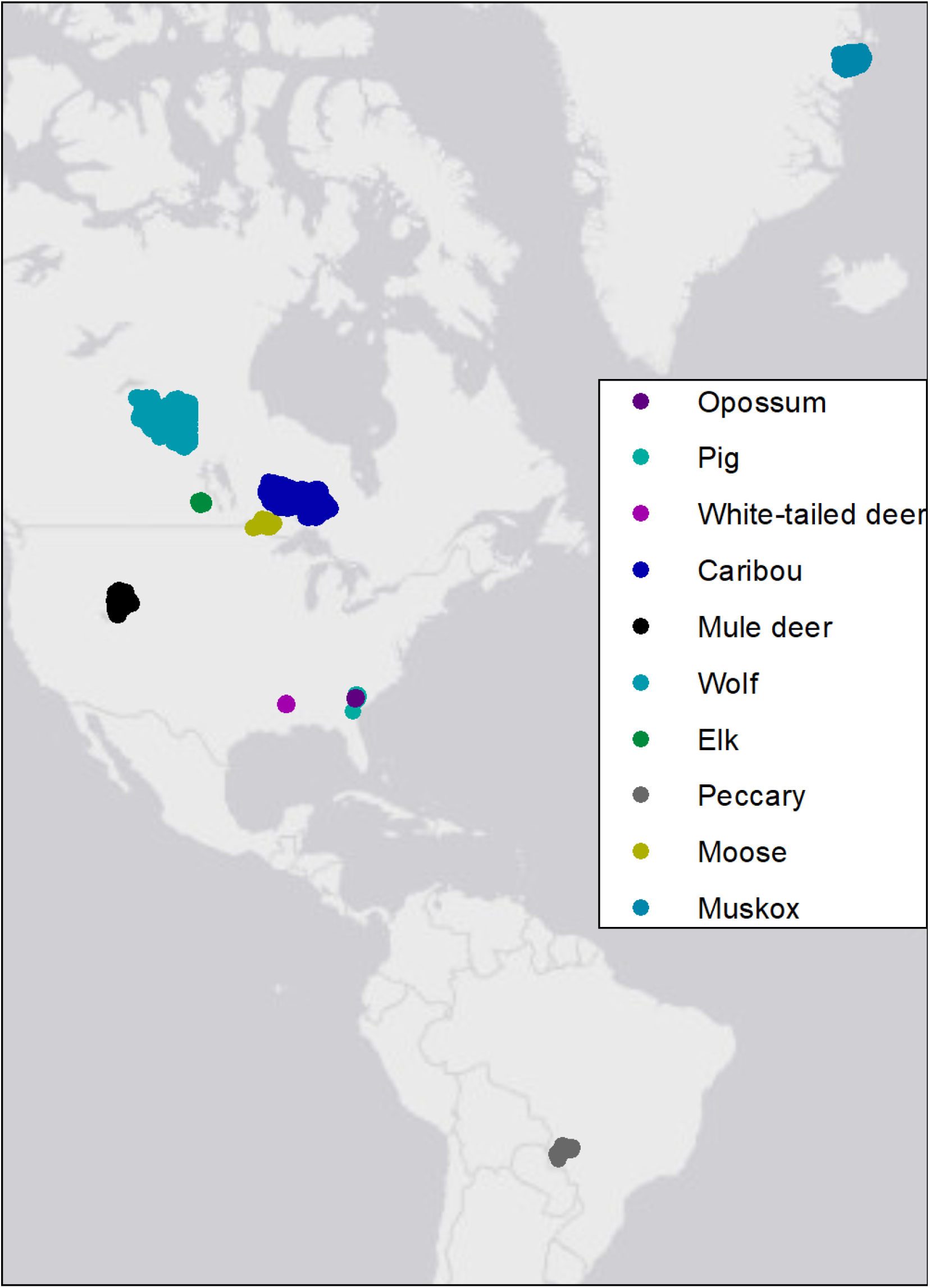
Data distribution. Geographic locations of GPS datasets (5,678,623 GPS relocations) across 511 individually collared members of 10 species.

We generated a 1:1 sample of availability (i.e. 1 available location per animal relocation) within each animal’s 99% home range as estimated using the function `kernelUD()` in R package `adehabitatHR` with the default bandwidth estimator. We extracted centered-and-scaled (numeric) and binary (categorical) landscape data to animal relocations and available locations and fit a RSF to each animal in each dataset using logistic regression (i.e. 511 individual models; Table S2). For simplicity, we used only linear main effects for each predictor in a given RSF; however, we emphasize that more complex effects (e.g. non-linear and interaction terms) may be identically investigated using the appropriate non-linear transformation or multiplicative product on the resource layer(s) prior to calculation. Note that, although our equations operate on a single resource layer at a time, our analysis uses RSFs with multiple layers. This procedure thus tests whether multiple layers may be analyzed one-at-a-time to ascertain the number of animals and fixes required to estimate the *β*-value for each layer.

### Empirical Validation: *M*

After fitting each RSF, we calculated the mean selection coefficient 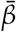 for each landscape layer across individuals within a species. Assuming 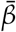 was an accurate estimate of population-level selection *β*, we asked: how many animals *M* were necessary to estimate *β*? We calculated Var[*R*(*X*_*β*_)] for each centered-and-scaled or binary raster within each animal’s 99% range according to Equation (4) and the resulting values of *N* according to Equation (3). We generated empirical distributions of 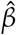 and 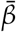 as described in Supplementary Appendix D for *M* ∈ {2, ..., 30} using the average *N* and Var[*R*(*X*_*β*_)] as population-level estimates of each. We computed the empirical 95% intervals at a given *M* (i.e. *α* = 0.05). The value of *M* at which the empirical interval no longer contains 0 is the predicted minimum *M* necessary to estimate *β* with 95% confidence, *M*_*pred*_ (i.e. the minimum integer *M*_0.05,0.05_(*β*); Equation (8)).

For comparison with observation, we then resampled the estimated selection coefficients for each individual within a species. For a given *M* ∈ {2, ..., 30} as above, we generated 4000 samples of *M*_*i*_ observed selection coefficients and calculated 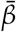 for each (i.e. 4000 mean selection coefficients assuming *M*_*i*_ animals). This represents the observed distribution of possible 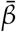 for *M*_*i*_ sampled animals, assuming the total pool of animals is a representative sample. Finally, for each *M* we calculated the grand mean 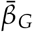 and the empirical 95% interval of 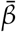. The value of *M* at which the empirical interval no longer contains 0 is the observed minimum *M* necessary to estimate *β* with 95% confidence, *M*_*obs*_, and should correspond to *M*_*pred*_

### Empirical Validation: *N*

The *M* validation procedure described above assumes that, on average, sufficient relocations *N* were available to estimate *M*. Now we consider: for a given individual-level selection coefficient *β*, do we have sufficient *N* to reject the null hypothesis for a given animal? We randomly sampled 1 animal from each dataset and calculated Var[*R*(*X*_*β*_)] within the animal’s 99% range using the animal’s specific RSF model coefficients as *β*. From this we calculated the predicted number of relocations *N*_*pred*_ necessary to estimate *β* given Var[*R*(*X*_*β*_)] (i.e. *N*_0.05,0.05_(*β*); Equation (3)).

For comparison, we resampled *N*_*sam*_ relocations with replacement from the animal’s dataset, where 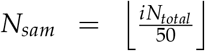, *i* ∈ {1, ..., 25}, and *N*_*total*_ is the total number of relocations recorded for that animal. This unconventional sequence was selected because (i) it produced a comparable number of observed values of *N* to that in the *M* validation procedure (25 observed *N* vs. 29 pairings of *M*_*pred*_ and *M*_*obs*_) while (ii) keeping the increments small enough to retain detail given that estimates of *N* can be orders of magnitude larger than those of *M*. We generated 4000 samples of *N*_*sam,i*_ relocations and fit an RSF to each individual sample (i.e. 4000 RSFs assuming *N*_*sam,i*_ relocations). We retained all originally generated available locations in each RSF so as to maintain a constant availability kernel between RSFs with different relocations. We then calculated the mean selection coefficient 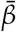 and its 95% empirical interval at a given *N*_*sam*_. The value of *N_sam_* at which the empirical interval no longer contains 0 is the observed minimum *N* necessary to reject *H*_0_ : *β* = 0 at significance level *p* ≤ 0.05, *N*_*obs*_, and should correspond to *N*_*pred*_.

## Results

The equations (3, 8) at the basis of our methods provide analytically predicted values for the number of relocations *N* and the number of animals *M* required to paramaterize an RSF. Simple 1-to-1 plots of *N*_*pred*_ vs. *N*_*obs*_ and *M*_*pred*_ vs. *M*_*obs*_ across all 10 species revealed remarkable agreement between observation and prediction (Fig. 3). Interestingly, 1 outlier was identified for *N* and 1 for *M*. Visual inspection of the data revealed that these outliers occurred alongside availability samples within individual RSFs that did not properly describe the true spatial integral of resource availability (i.e. ∫_Ω_*A*(**x′**)*W*(**x′**)*d***x′**; Equation (1)). That is, the 1:1 used/available sampling protocol undersampled the available space. Thus, *N*_*pred*_ and *M*_*pred*_ can be sensitive to insufficient spatial sampling of availability, and care should be taken to to avoid such undersampling before applying these methods.

**Figure 3:**
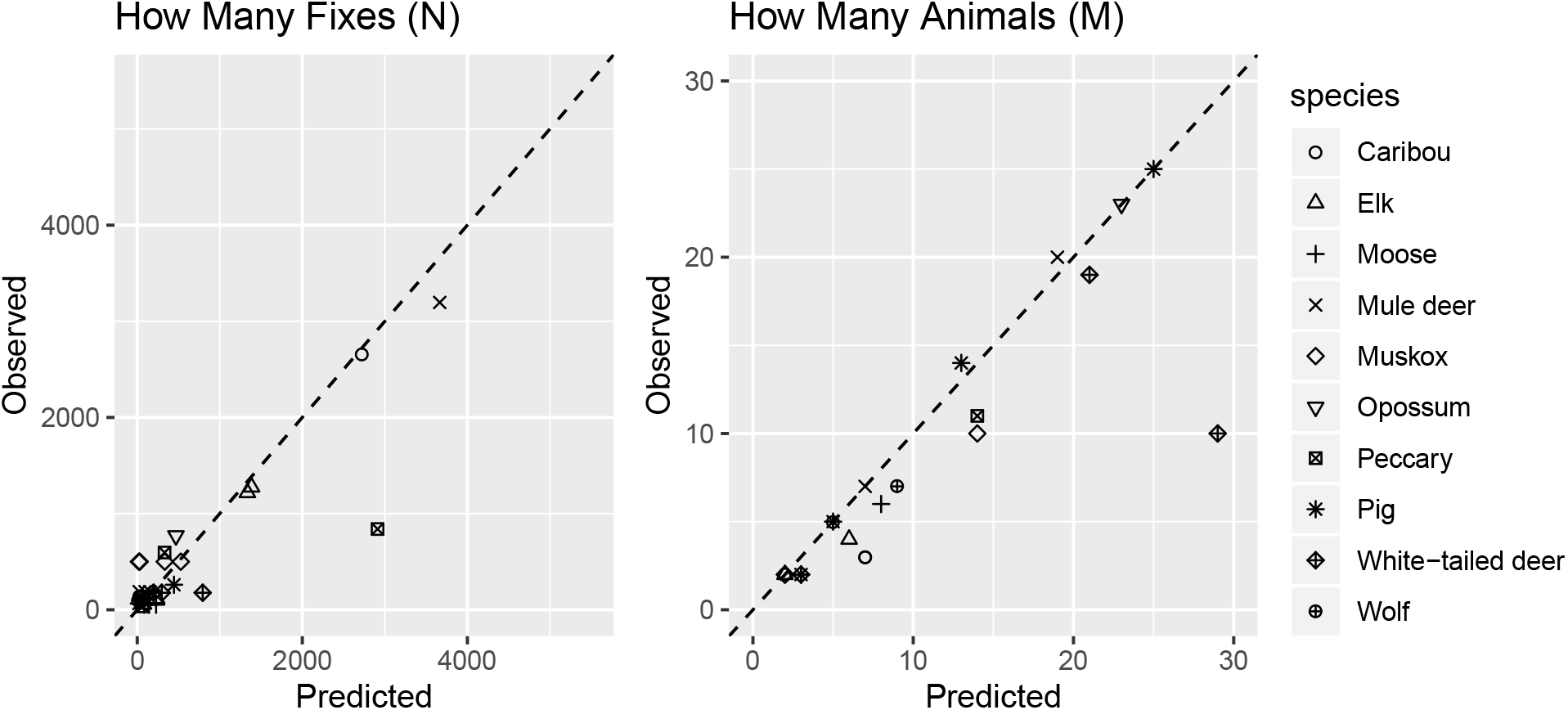
1-to-1 comparison of predicted and observed M and N. Three outliers are observed for *N* and one for *M* due to mismatch between sampled and true availability within the animals’ 99% ranges. Dashed lines are those with gradient 1 crossing through the origin.

Given this, we then asked, what is the role of the definition of availability (sensu Johnson 1980) in shaping these relationships? Our original calculations of *N*_*pred*_ and *M*_*pred*_ used individual availability (i.e. each animal has its own available resources within its unique 99% KDE). We repeated our calculations of *N*_*pred*_ and *M*_*pred*_, and bootstrap estimation of *N*_*obs*_ and *M*_*obs*_, using 2 additional availability definitions that varied the spatial extent of availability for a given animal: (i) within the entire collection of 99% KDEs (i.e. animals have access to resources within all KDEs equally), and (ii) within the entire site (i.e. animals have access to all resources within the study site, including those outside of 99% KDEs). This mimics the problem of sufficiently sampling availability described above, but now availability is driven by conceptual or ecological definitions rather than by the sampling protocol itself. Similar consistency in 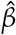 was observed across *M* within a given definition of availability, but the sign and magnitude of 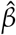 varied with availability from individual- to site-level (Fig. 4). Despite the change in sign and magnitude, Equation (8) is able to calculate *M*_*pred*_ consistent with observation across availability definitions. By inclusion, given that *N*_*pred*_ is a component of *M*_*pred*_ (see Equation (5)), we also observe that Equation (3) is consistent with observation across availability definitions.

**Figure 4:**
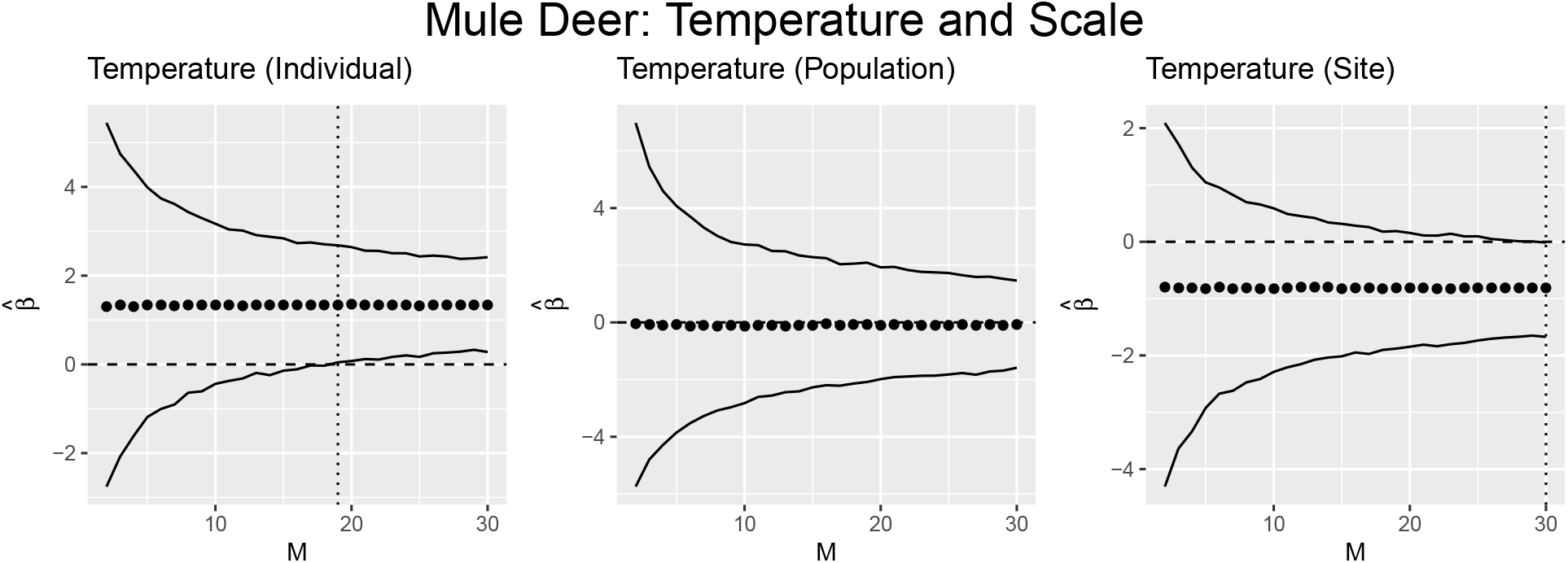
Comparison of predicted M across orders of availability. *M*_*pred*_ (vertical dotted line) changes depending on whether availability for the RSF is defined at the scale of the individual (each animal has its own available locations within its own 99% KDE), population (all animals have equal access to resources within all animal’s 99% KDEs), or site (all animals have equal access to resources across the entire site). If no vertical dotted line occurs, then *M*_*pred*_ > 30.

Lastly we asked, what are the primary drivers of *N*_*pred*_ and *M*_*pred*_ as estimated by Equations (3, 8)? A key outcome of our method is that this question can be answered analytically, by simply inspecting Equations (3, 8). Equation (3) shows that *N*_*pred*_ is inversely correlated to both Var[*R*(*X*_*β*_)] and *β*^2^, indicating that as either landscape variation or selection strength increase, so must *N*_*pred*_. Similarly, because *β*^2^ is contained in the denominator of Equation (8), *M*_*pred*_ must decrease with increasing selection strength. To demonstrate this graphically, we plotted log-log regressions of *N*_*pred*_ and *M*_*pred*_ against Var[*R*(*X*_*β*_)] and |*β*|, respectively, using data from all 10 species to evaluate whether these analytical predictions bear out under real data scenarios (Fig. 5). Per the analytical predictions, both *N*_*pred*_ and *M*_*pred*_ declined as their respective predictors (landscape variation or habitat selection strength) increased. It is also worth noting that inclusion of both predictors within the same log-log regression (i.e. *M*_*pred*_ as a function of both Var[*R*(*X*_*β*_)] and |*β*|) returned *R*^2^ = 1, as expected given that *N*_*pred*_ and *M*_*pred*_ are determined only by Var[*R*(*X*_*β*_)] and *β*.

**Figure 5:**
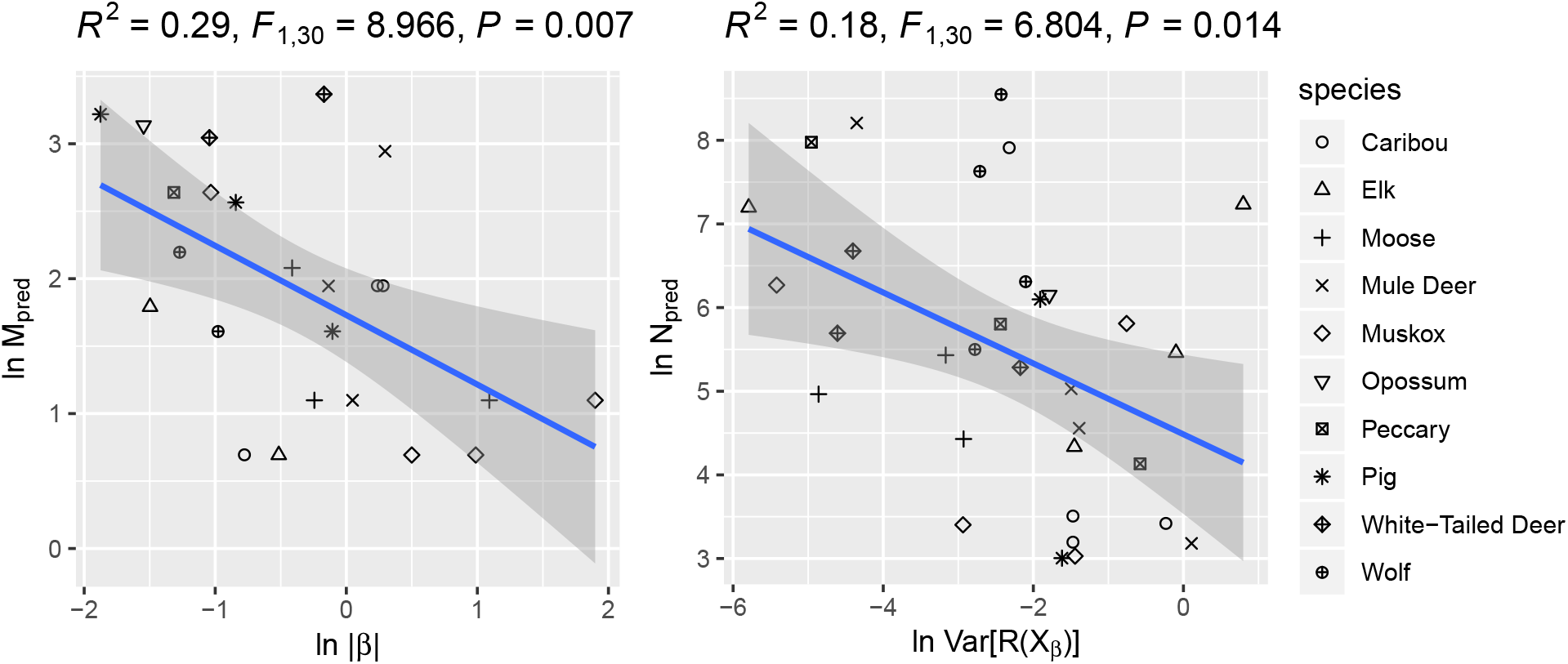
Log-log regressions of predicted M vs. |*β*| and predicted N vs. Var[*R*(*X*_*β*_)]. The predicted number of animals necessary *M*_*pred*_ declines with increasing absolute magnitude of selection (i.e. stronger effects require fewer animals to estimate), and the predicted number of relocations *N*_*pred*_ declines with increasing landscape complexity.

## Discussion

Conventional wisdom regarding sample size in RSA holds that a sample size of *M* ≥ 30 animals tagged is necessary for consistent and reliable inference (Leban *et al.* 2001; Hebblewhite & Haydon 2010). Convention also holds that more complex landscapes (i.e. those with higher landscape variance Var[*R*(*X*_*β*_)]) require more relocations per animal *N* to characterize selection (Wisz *et al.* 2008). Our analytical models and validation procedures return a contrasting set of results, contradicting conventional wisdom. First, we found that *M*_*pred*_ was often (but not always) substantially less than 30, and this prediction strongly agreed with observation based on resampling of GPS-based telemetry across a variety of ecologically contrasting species (Figs. 3, S1-S20). Strikingly, our analytical results show conclusively that *M* can only decline with increasing absolute magnitude of *β* (Equation (8)), indicating the most biologically relevant effects (i.e. those with the greatest |*β*|) can often be estimated with only a few animals (Fig. 5). This reveals important ethical and budgetary implications for wildlife studies. For example, consider the mule deer dataset containing 106 tagged individuals (Table S1). Our findings show that the strongest effects on the utilization distribution (i.e. selection for temperature, evergreen forest, and shrublands) may be estimated with fewer than 20 animals (Fig. S20), i.e. 80% fewer animals than were used. This means that, using a conservative estimate of US$2,450 for each GPS collar and data fees (K. L. Monteith, pers. obs.), if the sole aim of the study were to identify the relevant resource drivers of animal distributions as in typical RSF studies, this project would have overspent by $210,700 (excluding researcher/technician effort, which has significant cost in itself). Compared to the popular approach of tagging as many animals as possible and constructing phenomenological models to identify ecological mechanisms post hoc (colloquially referred to as “collar-and-foller”; Dunn 2004; Fieberg & Johnson 2015), our analytical results suggest researchers start with efforts aimed at constructing a priori hypotheses and associated models, then use our Equations (3, 8) to estimate the number of animals and locations per animal required for the study aims (Johnson *et al.* 2015).

Second, *N*_*pred*_ (the number of relocations per individual) also strongly agreed with observation, with both predicted and observed *N* in the 1000s or larger (Fig. 3). This agrees with findings that within-replicate sample sizes should generally be large (e.g. Wisz *et al.* 2008); however, our analytical expressions also conclusively demonstrate that *N* is directly calculable (Equation (3)) and as with *M* is expected to generally decline with increasing Var[*R*(*X*_*β*_)] and *β*. These conclusions for both *M* and *N* are not only analytically proven but are additionally supported by real data bearing out the analytical predictions (Figs. 3–5). As such, our findings demonstrate that not only are *M* and *N* imminently calculable given a known landscape and some expectation of *β*, but the expected trends in *M* and *N* with respect to landscape complexity and the strength of animal preference are precisely opposite those predicted by conventional wisdom and previous studies.

Why are our results contrary to so much of the preceding literature? One possibility could lie in the “golden rule” of sample size, i.e. that *M* ≥ 30 is required for a sample size sufficient to invoke the Central Limit Theorem and assume a roughly normal distribution of possible sample means (Aho 2014, p. 154), or to ignore non-normality because a model structure is somehow “robust” to non-normality (e.g. Hector 2015, p. 48). This is reinforced by an absence of mathematical attention to the sample size question. Previous studies have used simulation or empirical analyses to explore sample size sufficiency within particular species or systems (e.g. Leban *et al.* 2001; Loe *et al.* 2012; Sequeira *et al.* 2019), leading to conclusions that are quite specific to a given study but then are widely adopted as inferring pattern across all systems. By defining the problem mathematically (i.e. at what values of *M* and *N* do we reject the null hypothesis 100(1 − *α*)% of the time at significance *p*?), we instead arrive at general analytical solutions that then may be tested with simulations and empirical analyses that are specifically designed for those solutions, rather than relying on intuitive but incorrect assumptions about the relationships between landscape variation relative to selection strength and RSA sample size sufficiency.

Our calculations show that the required *M* and *N* for a given study are dependent entirely on |*β*| and Var[*R*(*X*_*β*_)]. The latter can be directly calculated given a landscape and an expectation for *β*, but selecting an appropriate expected *β* is a critical step in estimating *M* and *N*. For *a priori* planning this could be accomplished using expert knowledge and previous literature; however, there may be no conceivable prior expectation of *β* in some RSA exercises. In such a case, one may elect to perform for example a sensitivity analysis given a range of *β* to select conservative estimates of *M* and *N*. Further, observe that *β* is often affected by a variety of ecological phenomena, including resource availability, competitor density, and seasonal effects (Mysterud & Ims 1998; McLoughlin *et al.* 2010; van Beest *et al.* 2016; Paolini *et al.* 2018). This implies that Equations (3 & 8) estimating *N* and *M* respectively are in fact hierarchical with dependencies not only on landscape variance (i.e. Var[*R*(*X*_*β*_)]) but also landscape composition and structure as they determine *β*. In scenarios where we are uncertain about possible values of *β*, we may construct informed models suggesting likely values of *β* given an expectation for how the animal should behave as resource availability changes (e.g. generalized functional response models; Matthiopoulos *et al.* 2011). Such a hierarchical approach “borrows” information from the functional response model to provide a more ecologically informed range of possible *β* for a sensitivity analysis (Hobbs & Hooten 2015).

Our results also provide new insight into the importance of sufficient spatial sampling of availability. There was 1 outlier in the 1-to-1 comparison of *N*_*pred*_ and *N*_*obs*_, and 1 in that of *M*_*pred*_ and *M*_*obs*_ (Fig. 3). These occurred because the 99% range of the animals under observation was so large, and the underlying landscape rasters so finely grained, that our 1:1 use/availability sample did not accurately portray the spatial integral of availability ∫_Ω_*A*(**x′**)*W*(**x′**)*d***x′** (Equation (1)). This caused *M*_*pred*_ and *N*_*pred*_ to be based on a different, incomplete availability set compared to the fitted RSFs. This highlights an unexpected but critical conclusion: the sampling intensity for availability in RSF-styled models should be only as large as necessary to correctly characterize the availability integral. Previous RSF-styled studies (including SSF) have almost exclusively sampled availability as we did here using ratios (i.e. 1:1, 1:10, 1:100, etc.; e.g. Boyce & McDonald 1999; Fortin *et al.* 2005; Street *et al.* 2016). This encourages either sampling at an intensity insufficient to approximate the spatial integral (as occurred here for outlying points in Fig. 3), or at too great an intensity leading to overinflated sample sizes and biased standard errors, confidence intervals, and *p*-values. Both scenarios may affect inference, but despite these issues no general rule has been promoted for availability sampling in RSA. Based on our findings, we propose that this rule should be regular (non-random) sampling at a spatial interval equal to the resolution of the underlying landscape data such that every possible location within the availability boundary is considered. This would produce an availability observation for every raster pixel and thus overlap between used and available locations. Although it is suggested that such overlap is to be avoided (e.g. Wisz *et al.* 2008), logically a used location must also be available otherwise it cannot be selected, and removing used locations from availability can potentially omit important effects from the availability sample. Our equations indicate that this overlap is required by the mathematics of resource selection.

This finding reinforces that defining resource availability at the scale of the estimated model is a critical first step in planning a RSA. Our multi-scale analysis of mule deer produced remarkably different estimates for *M* at each of the three definitions of availability (site-wide, population-wide, and individual availability; Fig. 4), indicating that failure to properly define the available space can lead to incorrect estimates of both *M* and *N*. This is not a new finding; the importance of properly defining what is available for an animal to select is a long-standing issue in RSA research (e.g. Johnson 1980; Boyce & McDonald 1999; Fortin *et al.* 2005). However, the difficulty of calculating *M* and *N* for planning a RSA study increases with the biological scale of the intended model. Site-wide availability assumes all animals have access to resources on the entire landscape and is similar in concept to first-order selection (i.e. where the species is located; Johnson 1980), but availability may be sampled as a regular grid across the entire site. Population-wide availability refines the scale toward second-order selection (i.e. where animals situate their home ranges), but accurately defining a perimeter for the likely population range *a priori* within which to sample availability is non-trivial. This becomes even more difficult under individual availability; how can we anticipate the size and placement of individual home ranges? A feasible solution may be to delineate population boundaries and within this delineation generate random ranges with area determined by the literature and expert knowledge. This would enable calculation of an average theoretical availability for any animal in the study site with appropriate standard error. This could then be used to produce an average prediction for *M* and *N*, and associated confidence limits, across the average home range composition.

We approached this analysis with the specific intention of evaluating how many GPS-tagged animals *M* are needed for RSF estimation, but there are many RSF applications that do not seek *M* or require GPS-tagging (e.g. plant distributions). For example, RSAs estimated for rare species will typically lack sufficient data for individual-based estimation of the utilization distribution *u*(*x*) such that *M* is irrelevant and only *N* need be evaluated. RSAs can be sensitive to small sample sizes (Wisz *et al.* 2008), yet they often generate accurate predictions for rare species with small datasets (McCune 2016), suggesting that for some rare species smaller *N* is sufficient to achieve a robust model. Our findings permit evaluation of this. Consider a hypothetical scenario where RSA is conducted for a rare species with 100 observations and *β* is recorded. Here, Equations (3–4) could be used to calculate *N*_*pred*_ as a *post hoc* metric of confidence assuming *β* is the true population/species-level average selection coefficient. If *N*_*pred*_ ≤ 100, then one could trust the outcome of the RSA; conversely, *N_pred_ >* 100 would indicate additional data collection is necessary. Where that is not possible, one could systematically adjust *z*_*α*_ and *z*_*p*/2_ (Equation (3)) to determine the percent confidence interval that rejects the null hypothesis *H*_0_ : *β* = 0 and establish a degree of confidence for model outcomes. Although there are issues with this approach (e.g. individual variation is ignored), this is a limitation of small datasets and not the equations identified here. Similarly, although we performed validation using GPS-based datasets, Equation (3) is agnostic to how data are collected and may also be applied to sessile organisms. Provided we can plausibly accept that *β* is roughly true and individual variation is either minimal or accommodated by the population-level *β* (presumably what has been estimated), our equations may be easily extended to evaluate most any RSA-based study.

We must emphasize that although *M* may only decline with increasing *β*, Equation (3) allows for a turning point to occur such that *N* initially decreases with |*β*| but eventually increases at very large |*β*| (see Supplemental Information, Equation (25)). When selection strength is particularly strong, smaller sample sizes make it much more likely to obtain perfect separation between used and unused resources. In such a case one must collect more data to observe the animal not using a resource unit it should strongly prefer (or in the case of negative selection, to observe it using a resource it should strongly avoid). Practically, this means that sampling intensity for RSA is a greater concern for specialist organisms than generalists because specialists should exhibit typically larger |*β*| for preferred/avoided resource units than generalists. Although the equations identified here allow us to directly calculate *N* for any landscape and expected selection strength, we should generally expect that specialists will require larger *N* for precise RSA estimation.

The equations identified here explicitly evaluate the compatibility of a dataset with a given hypothetical model (i.e. *β*). Calculating their solutions across gradients of *N* and *M* reveals how the number of data points (relocations) and number of replicates (animals) affect determination of compatibility. Rather than the values of *N* and *M* required to achieve statistical significance, we instead suggest these be used to determine the relevant sample sizes necessary to achieve “consistent” results, i.e. if we increase sampling intensity would we see substantial change in estimated coefficients? From this perspective, we conclude that the number of animals *M* required to consistently estimate the most biologically relevant effects in an RSA can be well below commonly touted sample size thresholds (i.e. *M* ≥ 30), particularly when selection strength is strong (Fig. 3, 5). Moreover, the number of required relocations *N* can also be quite small but tends toward larger sample sizes when landscape variation is small. The sufficiency of samples sizes *M* and *N* is dependent entirely on the strength of selection (|*β*|) and landscape variation with respect to selection strength (Var[*R*(*X*_*β*_)]). Rather than simply reporting sample sizes in RSA studies, researchers should pay explicit attention to the effect their sample size has on their findings. Regardless of study organism, ecosystem, or scenario, our equations may be equally applied to *any* RSF-based study to evaluate the consistency of expected outcomes given a dataset of a particular size. This will partially address the so-called “replicability crisis” by explicitly characterizing the consistency of model outputs in relation to sample sizes and effect sizes, thereby increasing reader (and reviewer) confidence in such studies. Similarly, editors and reviewers should abandon preconceived notions of what makes a sufficient sample size in RSA in favor of evaluating the sensitivity of findings to sample size based on the mathematical rules identified here, for it is also feasible (and indeed demonstrable) that consistent findings can be achieved with as few as *N* = 100 relocations per animal and *M* = 2 animals (Fig. 3). Because *M* and *N* can be easily calculated provided knowledge of ecological and landscape effects, we argue that such calculations should henceforth be a mandatory component for all RSA studies.

## Supporting information

Finding Sums of Squares R Supplement

## Acknowledgments

We thank the Movement Ecology Special Interest Group of The British Ecological Society for valuable discussion regarding this topic, in particular Marie Auger-Métheé. GMS thanks the Mississippi Agricultural and Forestry Experiment Station (MAFES); the Forest and Wildlife Research Center (FWRC); the United States Department of Agriculture National Institute of Food and Agriculture (USDA NIFA); and the Mississippi Department of Wildlife, Fisheries, and Parks (MDWFP) for supporting this research and associated data collection. JRP thanks the School of Mathematics and Statistics at the University of Sheffield for granting him study leave which has helped enable the research presented here. CMP, LJN, and EV respectfully acknowledge that Riding Mountain National Park is the traditional homeland of the Anishinabe People and the Métis Nation, within Treaty 2 territory and at the crossroads of Treaties 1 and 4. Contributions of SD and BKS were partially supported by the Mississippi State University Extension Service (MSUES), FWRC, and MDWFP. Contributions of JCB, OER, PES, GD, DAK, and DAB were partially supported by USDA Animal and Plant Health Inspection Service (APHIS), Wildlife Services (WS), National Wildlife Research Center (NWRC), and U.S. Department of Energy (DOE) through Cooperative Agreement number DE-FC09-07SR22506 with the University of Georgia Research Foundation. Contributions of ARR and JMF were supported by the Ontario Ministry of Natural Resources and Forestry (OMNRF). Contributions of KLM and SPD were supported by Wyoming Game and Fish Department (WGFD), Bureau of Land Management (BLM), Muley Fanatic Foundation, Boone and Crockett Club, Wyoming Wildlife and Natural Resources Trust, Knobloch Family Foundation, Wyoming Animal Damage Management Board, Wyoming Governor’s Big Game License Coalition, Bowhunters of Wyoming, Wyoming Outfitters and Guides Association, United States Forest Service (USFS), and United States Fish and Wildlife Service (USFWS). Contributions of CMP, LJN, and EV were supported primarily by Parks Canada Agency (Riding Mountain National Park of Canada) and the Natural Science and Engineering Research Council of Canada (NSERC). Contributions of FMvB, LTB, and NMS were supported by the AUFF Starting Grant (AUFF-F-2016-FLS-8-16).

## Author Contributions

GMS conceived and directed the project and developed the validation and resampling framework. JRP derived the analytic expressions, and GMS and JRP conducted the data analyses in collaboration with LB. GMS, JRP, and LB wrote the manuscript. Remaining authors collected and contributed data and contributed equally to edits and revisions.

## Data Accessibility

RSF and landscape data will be uploaded to a permanent repository following formal acceptance of this manuscript for publication.

