## Supplemental Appendices for "Solving the Sample Size Problem for Resource Selection Analysis"

### Supplementary Appendix A

The aim of this Appendix is to prove Equation (3) of the Main Text. We do this by analysing the likelihood function for the model given in Equation (2) of the Main Text. To this end, suppose $\mathbf{x}_1, \ldots, \mathbf{x}_N$ are independent datapoints. Suppose our model for the distribution of these data has the following probability density function (which is Equation 2 in the Main Text)

$$u(\mathbf{x}|\beta) = \frac{\exp[\beta R(\mathbf{x})]}{\int_\Omega \exp[\beta R(\mathbf{x}')] d\mathbf{x}'}, \tag{1}$$

where $R : \Omega \to \mathbb{R}$ is a bounded, integrable function. Then the likelihood of $\beta$, given the data $\mathbf{x}_1, \ldots, \mathbf{x}_N$, is

$$L(\beta|\mathbf{x}_1, \ldots, \mathbf{x}_N) = \frac{\exp\left[\beta \sum_{j=1}^{N} R(\mathbf{x}_j)\right]}{\left(\int_\Omega \exp\left[\beta R(\mathbf{x})\right] \mathrm{d}\mathbf{x}\right)^N}. \tag{2}$$

We begin by proving Proposition A.1, which gives a formula for the critical point of this likelihood function, together with a proof that it exists and is unique. Then Proposition A.2 shows that this unique critical point is a maximum. (It is worth noting that the existence and uniqueness of the MLE is already known in a more general setting (McCullagh, 1980), but proved here in our specific case, for completeness, and so it is clear where the formula for the MLE comes from.)

**Proposition A.1.** The critical points of the likelihood function from Equation (2) occur at the points where the following equation is solved

$$\frac{\int_\Omega R(\mathbf{x}) \exp[\beta R(\mathbf{x})]\mathrm{d}x}{\int_\Omega \exp[\beta R(\mathbf{x})]\mathrm{d}x} = \frac{1}{N}\sum_{j=1}^{N} R(\mathbf{x}_j). \tag{3}$$

Furthermore, there is a unique solution to Equation (3).

*Proof.* Differentiating Equation (2) with respect to $\beta$ gives

$$\frac{\partial L}{\partial \beta} = \exp\left[\beta \sum_{j=1}^{N} R(\mathbf{x}_j)\right] \frac{\int_\Omega \exp[\beta R(\mathbf{x})]\mathrm{d}\mathbf{x} \sum_{j=1}^{N} R(\mathbf{x}_j) - N \int_\Omega R(\mathbf{x}) \exp[\beta R(\mathbf{x})]\mathrm{d}\mathbf{x}}{\left[\int_\Omega \exp[\beta R(\mathbf{x})]\mathrm{d}\mathbf{x}\right]^{N+1}}. \quad (4)$$

This is equal to zero precisely when the numerator of the fraction in Equation (4) is zero, i.e. when

$$\int_\Omega \exp[\beta R(\mathbf{x})]\mathrm{d}\mathbf{x} \sum_{j=1}^{N} R(\mathbf{x}_j) - N \int_\Omega R(\mathbf{x}) \exp[\beta R(\mathbf{x})]\mathrm{d}\mathbf{x} = 0. \quad (5)$$

Equation (5) rearranges to give Equation (3), so we have shown that the critical points of the likelihood function occur precisely when Equation (3) is satisfied.

We now need to show that the solution to Equation (3) is unique. For this, we let $\epsilon > 0$ be arbitrarily small and define $\tilde{R}(\mathbf{x}) = R(\mathbf{x}) - R_0$ where $R_0 = \max\{R(\mathbf{x})\} + \epsilon$. Then $\tilde{R}(\mathbf{x}) < 0$ for all $\mathbf{x}$. Let $\tilde{g}(\beta) = \int_\Omega \exp[\beta \tilde{R}(\mathbf{x})]\mathrm{d}\mathbf{x}$. Then $\tilde{g}'(\beta) = \int_\Omega \tilde{R}(\mathbf{x}) \exp[\beta \tilde{R}(\mathbf{x})]\mathrm{d}\mathbf{x} < 0$, so $\tilde{g}(\beta)$ is a decreasing function of $\beta$.

Also, $\tilde{g}''(\beta) = \int_\Omega \tilde{R}^2(\mathbf{x}) \exp[\beta \tilde{R}(\mathbf{x})]\mathrm{d}\mathbf{x}$, which is greater than 0, since $R^2(\mathbf{x}) \geq 0$ and $\exp[\beta R(\mathbf{x})] > 0$. Therefore $\tilde{g}'(\beta)$ is an increasing function of $\beta$. Consequently, $\tilde{g}'(\beta)/\tilde{g}(\beta)$ is an increasing function of $\beta$, so the following has a unique solution

$$\frac{\tilde{g}'(\beta)}{\tilde{g}(\beta)} = \frac{1}{N} \sum_{j=1}^{N} \tilde{R}(\mathbf{x}_j). \quad (6)$$

Let $g(\beta) = \int_\Omega \exp[\beta R(\mathbf{x})]\mathrm{d}\mathbf{x}$. Then Equation (3) can be written as

$$\frac{g'(\beta)}{g(\beta)} = \frac{1}{N} \sum_{j=1}^{N} R(\mathbf{x}_j). \quad (7)$$

Now suppose that $\beta^*$ is a solution to Equation (6). Then

$$\begin{aligned}\frac{g'(\beta^*)}{g(\beta^*)} &= \frac{\int_\Omega [\tilde{R}+R_0]\exp(\beta^* R_0)\exp(\beta^*\tilde{R}(\mathbf{x}))\mathrm{d}\mathbf{x}}{\int_\Omega \exp(\beta^* R_0)\exp(\beta^*\tilde{R}(\mathbf{x}))\mathrm{d}\mathbf{x}} \\ &= \frac{\int_\Omega \tilde{R}\exp(\beta^*\tilde{R}(\mathbf{x}))\mathrm{d}\mathbf{x}}{\int_\Omega \exp(\beta^*\tilde{R}(\mathbf{x}))\mathrm{d}\mathbf{x}} + \frac{R_0\int_\Omega \exp(\beta^*\tilde{R}(\mathbf{x}))\mathrm{d}\mathbf{x}}{\int_\Omega \exp(\beta^*\tilde{R}(\mathbf{x}))\mathrm{d}\mathbf{x}} \\ &= \frac{1}{N}\sum_{j=1}^{N}\tilde{R}(\mathbf{x}_j) + R_0 \\ &= \frac{1}{N}\sum_{j=1}^{N}[\tilde{R}(\mathbf{x}_j) + R_0] \\ &= \frac{1}{N}\sum_{j=1}^{N}R(\mathbf{x}_j), \end{aligned} \tag{8}$$

so $\beta^*$ is a solution to Equation (7), and hence Equation (3). By an identical argument, any solution to Equation (3) is also a solution to Equation (6). Therefore the values of $\beta$ that solve Equation (6) are exactly the same as those that solve Equation (7). Since there is a unique solution to Equation (6), it follows that the solution to Equation (3) is unique. □

**Proposition A.2.** The derivative of the likelihood function from Equation (2) has the following properties

1. $\lim_{\beta\to-\infty} L'(\beta) \geq 0$,
2. $\lim_{\beta\to\infty} L'(\beta) \leq 0$.

*Proof.* From Equation (4), we see that the sign of $L'(\beta)$ is the same as the sign of

$$\int_\Omega \exp[\beta R(\mathbf{x})]\mathrm{d}\mathbf{x}\sum_{j=1}^{N}R(\mathbf{x}_j) - N\frac{\int_\Omega R(\mathbf{x})\exp[\beta R(\mathbf{x})]\mathrm{d}\mathbf{x}}{\int_\Omega \exp[\beta R(\mathbf{x})]\mathrm{d}\mathbf{x}}. \tag{9}$$

Now,

$$\lim_{\beta\to-\infty}\int_\Omega R(\mathbf{x})\frac{\exp[\beta R(\mathbf{x})]}{\int_\Omega \exp[\beta R(\mathbf{x}')]\mathrm{d}\mathbf{x}'}\mathrm{d}\mathbf{x} = \delta(x - \mathrm{argmin}_{\mathbf{x}}\{R(\mathbf{x})\}), \tag{10}$$

where $\delta(\mathbf{x})$ is the Kronecker delta function. Therefore

$$\lim_{\beta\to-\infty}\left\{\frac{\int_\Omega R(\mathbf{x})\exp[\beta R(\mathbf{x})]\mathrm{d}\mathbf{x}}{\int_\Omega \exp[\beta R(\mathbf{x})]\mathrm{d}\mathbf{x}}\right\} = \int_\Omega \delta(\mathbf{x}-\mathrm{argmin}_{\mathbf{x}}\{R(\mathbf{x})\})R(\mathbf{x})\mathrm{d}\mathbf{x}$$
$$= \min\{R(\mathbf{x})\} \tag{11}$$

It follows that

$$\lim_{\beta\to-\infty}\left\{\int_\Omega \exp[\beta R(\mathbf{x})]\mathrm{d}\mathbf{x}\sum_{j=1}^{N} R(\mathbf{x}_j) - N\frac{\int_\Omega R(\mathbf{x})\exp[\beta R(\mathbf{x})]\mathrm{d}\mathbf{x}}{\int_\Omega \exp[\beta R(\mathbf{x})]\mathrm{d}\mathbf{x}}\right\}$$
$$= \sum_{j=1}^{N} R(\mathbf{x}_j) - N\min\{R(\mathbf{x})\}. \tag{12}$$

Since $R(\mathbf{x}_j) \geq \min\{R(\mathbf{x})\}$, it follows that $\lim_{\beta\to-\infty} L'(\beta) \geq 0$. The argument showing $\lim_{\beta\to\infty} L'(\beta) \leq 0$ is analogous. □

Calculating $N_{\alpha,p}(\beta)$ from the Main Text requires an understanding of the standard error, $\sigma$, of our maximum likelihood estimator. A standard result relating the standard error to the Hessian of the log-likelihood function (Andersen, 1970) implies that, in our case, $\sigma \approx \sqrt{[-l''(\beta|\mathbf{x}_1,\ldots,\mathbf{x}_N)]^{-1}}$, where $l(\beta|\mathbf{x}_1,\ldots,\mathbf{x}_N) = \ln[L(\beta|\mathbf{x}_1,\ldots,\mathbf{x}_N)]$. This becomes exact as $N\to\infty$.

Therefore, to calculate $\sigma$, we begin by taking the natural logarithm of Equation (2), to give

$$l(\beta|\mathbf{x}_1,\ldots,\mathbf{x}_N) = \ln[L(\beta|\mathbf{x}_1,\ldots,\mathbf{x}_N)] = \beta\sum_{j=1}^{N} R(\mathbf{x}_j) - N\ln\left(\int_\Omega \exp\left[\beta R(\mathbf{x})\right]\mathrm{d}\mathbf{x}\right). \tag{13}$$

The first and second derivatives of Equation (13) are, respectively,

$$l'(\beta|\mathbf{x}_1,\ldots,\mathbf{x}_N) = \sum_{j=1}^{N} R(\mathbf{x}_j) - N\frac{\int_\Omega R(\mathbf{x})\exp\left[\beta R(\mathbf{x})\right]\mathrm{d}x}{\int_\Omega \exp\left[\beta R(\mathbf{x})\right]\mathrm{d}x},$$
$$l''(\beta|\mathbf{x}_1,\ldots,\mathbf{x}_N) = -N\frac{\int_\Omega \exp\left[\beta R(\mathbf{x})\right]\mathrm{d}x\int_\Omega R^2(x)\exp\left[\beta R(\mathbf{x})\right]\mathrm{d}x - \left(\int_\Omega R(\mathbf{x})\exp\left[\beta R(\mathbf{x})\right]\mathrm{d}x\right)^2}{\left(\int_\Omega \exp\left[\beta R(\mathbf{x})\right]\mathrm{d}x\right)^2}. \tag{14}$$

Consequently,

$$\sigma \approx \frac{1}{\sqrt{N \mathrm{Var}[R(X_\beta)]}}, \tag{15}$$

where

$$\mathrm{Var}[R(X_\beta)] = \frac{\int_\Omega R^2(\mathbf{x}) \exp\left[\beta R(\mathbf{x})\right] \mathrm{d}\mathbf{x}}{\int_\Omega \exp\left[\beta R(\mathbf{x})\right] \mathrm{d}\mathbf{x}} - \left( \frac{\int_\Omega R(\mathbf{x}) \exp\left[\beta R(\mathbf{x})\right] \mathrm{d}\mathbf{x}}{\int_\Omega \exp\left[\beta R(\mathbf{x})\right] \mathrm{d}\mathbf{x}} \right)^2. \tag{16}$$

Notice that $\mathrm{Var}[R(X_\beta)]$ is the variance of $R(X_\beta)$, where $X_\beta$ is a random variable with the probability density function given by Equation (1).

Recall that $N_{\alpha,p}(\beta)$ was defined to be the minimum number of samples, $N$, required so that we expect to reject the null hypothesis of $\beta = 0$ (vs. alternative $\beta \neq 0$), at a significance level of $p$, in $100(1-\alpha)\%$ of experiments. In other words, it is the minimum $N$ such that $P(|\hat{\beta}| > \sigma z_{p/2}) \geq 1 - \alpha$, where $\hat{\beta}$ is the maximum likelihood estimator for $\beta$.

Assuming that $\hat{\beta} \sim N(\beta, \sigma^2)$ (i.e. $\hat{\beta}$ is normally distributed with mean $\beta$ and variance $\sigma^2$), the symmetry of the normal distribution means that

$$P(\hat{\beta} > \sigma z_{p/2}) + P(\hat{\beta} < -\sigma z_{p/2}) \geq 1 - \alpha. \tag{17}$$

Assume, without loss of generality, that $\beta > 0$. Then, for any typically-used $p$-value (i.e. $p \leq 0.05$), we have that $P(\hat{\beta} < -\sigma z_{p/2})$ is much less than $P(\hat{\beta} > \sigma z_{p/2})$, giving the following approximate inequality

$$P(\hat{\beta} > \sigma z_{p/2}) \gtrsim 1 - \alpha, \tag{18}$$

which rearranges to give

$$P\left( \frac{\hat{\beta} - \beta}{\sigma} > z_{p/2} - \frac{\beta}{\sigma} \right) \gtrsim 1 - \alpha. \tag{19}$$

By the assumption that $\hat{\beta} \sim N(\beta, \sigma)$, Equation (19) rearranges to give

$$1 - \Phi\left( z_{p/2} - \frac{\beta}{\sigma} \right) \gtrsim 1 - \alpha, \tag{20}$$

where $\Phi(\cdot)$ is the cumulative distribution function of the standard normal distribution. Then

Equation (20) becomes

$$-\left(z_{p/2}-\frac{\beta}{\sigma}\right)\gtrsim -\Phi^{-1}(\alpha)=z_\alpha, \tag{21}$$

where $z_\alpha=\Phi^{-1}(1-\alpha)=-\Phi^{-1}(\alpha)$. Equation (21) rearranges to give the following inequality for $\beta$

$$\beta\gtrsim\sigma(z_\alpha+z_{p/2})=\frac{z_\alpha+z_{p/2}}{\sqrt{N\mathrm{Var}[R(X_\beta)]}}, \tag{22}$$

where the equality in (22) comes from Equation (15). Finally, (22) rearranges to give

$$N\gtrsim\frac{(z_\alpha+z_{p/2})^2}{\mathrm{Var}[R(X_\beta)]}\beta^{-2}, \tag{23}$$

from whence the value of $N_{\alpha,p}(\beta)$ in Equation (3) of the Main Text arises.

The derivative of $N_{\alpha,p}(\beta)$ is given as follows

$$\frac{\partial N}{\partial\beta}=-\frac{(z_\alpha+z_{p/2})^2}{\beta^3\mathrm{Var}[R(X_\beta)]^2}\left(2\mathrm{Var}[R(X_\beta)]+\beta\frac{\partial}{\partial\beta}\mathrm{Var}[R(X_\beta)]\right). \tag{24}$$

Define $F_n(\beta)=\int_\Omega R^n(\mathbf{x})\exp[\beta R(\mathbf{x})]\,\mathrm{d}\mathbf{x}$. Then Equation (24) is equal to zero (i.e. there is a turning point) when

$$0=2F_2(\beta)F_0^2(\beta)-2F_1^2(\beta)F_0(\beta)-\beta[F_0^2(\beta)F_3(\beta)-3F_0(\beta)F_1(\beta)F_2(\beta)+2F_1^3(\beta)]. \tag{25}$$

### Supplementary Appendix B

In this section, we explain how we calculated empirical values for $N_{\alpha,p}(\beta)$ from simulated data, as displayed in Fig. 1b of the Main Text. We used a simulated landscape, described in the Methods section of the Main Text and displayed in Fig. 1a from the Main Text. We constructed simulated datasets of 15 'virtual species', each of which has a space use distribution described by Equation (1) for a unique value of $\beta\in\{0.1,0.2,\dots,1.5\}$.

Constructing a simulated dataset of size $N$, for a virtual species with a given $\beta$-value, requires taking $N$ independent samples from the distribution in Equation (1). For each pair of values $(\beta,N)$ that we tested, we constructed 1,000 simulated datasets. For each dataset, we used conditional logistic regression (using the `clogit()` function in `R` from the `survival` package), with 100 controls per case, to test the null hypothesis $\beta=0$ against the alternative $\beta\neq0$ at a $p=0.05$ significance level. We then calculated, for each pair $(\beta,N)$, the proportion of datasets for which the null hypothesis was rejected. We denote this proportion by $q(\beta, N)$.

For each $\beta$, $q(\beta, N)$ is a roughly increasing function of $N$, possibly with a small amount of noise owing to the random sampling. By smoothing $q(\beta, N)$ if necessary (to remove the noise), we therefore find a unique value, $N_{0.5,0.05}(\beta)$, for which $q(\beta, N) \geq 0.5$ if and only if $N \geq N_{0.5,0.05}$. Similarly, $N_{0.05,0.05}(\beta)$ is the unique value of $N$ for which $q(\beta, N) \geq 0.95$ if and only if $N \geq N_{0.05,0.05}$. These can be compared directly to the corresponding analytic values, with the same names, defined by Equation (3) of the Main Text. The result is displayed in Fig. 1b of the Main Text.

We also re-did the analysis using logistic regression (instead of conditional logistic regression). The method was the same except, instead of `clogit()`, we used the `glm()` function with an error distribution given by `family=binomial(link=logit)`. Resulting values for $N_{0.5,0.05}$ and $N_{0.05,0.05}$ were essentially identical (i.e. the majority were exactly the same integer values and the others were so close as to be almost impossible to distinguish by eye when superimposed on Fig. 1b of the Main Text).

### Supplementary Appendix C

In this section, we derive Equations (6) and (7) from the Main Text. Recall that $\beta_1, \ldots, \beta_M \sim N(\beta, s^2)$ are independent draws from a normal distribution with mean $\beta$ and variance $s^2$. Recall also that $\hat{\beta}_i \sim N(\beta_i, \sigma_i^2)$ where $\sigma_i$ is calculated from the number of locations gathered for animal $i$, using Equation (15), and that $\hat{\beta}$ is defined to be the mean of $\hat{\beta}_1, \ldots, \hat{\beta}_M$, so

$$\hat{\beta} = \frac{1}{M}\sum_{i=1}^{M}\hat{\beta}_i. \tag{26}$$

Since $\hat{\beta}_i - \beta_i \sim N(0, \sigma_i^2)$ and $\beta_i - \beta \sim N(0, s^2)$, we have that

$$\begin{aligned}
\hat{\beta} - \beta &= \frac{1}{M}\sum_{i=1}^{M}(\hat{\beta}_i - \beta) \\
&= \frac{1}{M}\sum_{i=1}^{M}(\hat{\beta}_i - \beta_i) + \frac{1}{M}\sum_{i=1}^{M}(\beta_i - \beta) \\
&\sim N\left(0, \frac{1}{M^2}\sum_{i=1}^{M}\sigma_i^2 + \frac{s^2}{M^2}\right),
\end{aligned} \tag{27}$$

from which Equation (6) in the Main Text follows.

To show Equation (7), define the sample variance as follows

$$V = \frac{1}{M}\sum_{i=1}^{M}(\hat{\beta}_i - \hat{\beta})^2. \tag{28}$$

We then have the following sequence of equalities to calculate the expectation of $V$

$$\begin{aligned} E[V] &= \frac{1}{M}\sum_{i=1}^{M}(\hat{\beta}_i - \hat{\beta})^2 \\ &= \frac{M-1}{M^2}\sum_{i=1}^{M} E[\hat{\beta}_i^2] - \frac{2}{M}\sum_{i=1}^{M}\sum_{j>i} E[\hat{\beta}_i\hat{\beta}_j] \\ &= \frac{M-1}{M^2}\sum_{i=1}^{M} E[\hat{\beta}_i^2] - \frac{2}{M}\sum_{i=1}^{M}\sum_{j>i} \beta_i\beta_j \\ &= \frac{M-1}{M^2}\sum_{i=1}^{M}(\sigma_i^2 + \beta_i^2) - \frac{2}{M}\sum_{i=1}^{M}\sum_{j>i} \beta_i\beta_j \\ &= \frac{M-1}{M^2}\sum_{i=1}^{M}\sigma_i^2 + \frac{1}{M}\sum_{i=1}^{M}\left(\beta_i - \frac{1}{M}\sum_{j=1}^{M}\beta_j\right)^2, \end{aligned} \tag{29}$$

where the third equality uses the fact that $\hat{\beta}_i$ and $\hat{\beta}_j$ are independent for $i \neq j$. Since $\beta_i \sim N(\beta, s^2)$ are also random variables, a direct calculation reveals

$$E[E(V)] = \frac{M-1}{M^2}\sum_{i=1}^{M}(\sigma_i^2 + s^2). \tag{30}$$

Thus $V$ is an unbiased estimator of the right-hand side of Equation (30). Rearranging Equation (30) gives the unbiased estimator for $s^2$ reported in Equation (7) of the Main Text.

### Supplementary Appendix D

In this section, we investigate numerically the effect of various parameter choices on the precision of our estimator for $s^2$ given in Equations (7) of the Main Text. At the same time, we numerically verify the validity our estimators for both $s^2$ and $\beta$ (the latter being given on Equation 6 of the Main Text).

To this end, we choose a variety of values of $\beta$, $s^2$, $M$, and $\sigma_1, \ldots, \sigma_M$. For each set of parameter values, we sample $\beta_1, \ldots, \beta_M \sim N(\beta, s^2)$, then $\hat{\beta}_i \sim N(\beta_i, \sigma_i^2)$ and calculate $\hat{\beta}$ and $\hat{s^2}$. We perform this sampling procedure 10,000 times to give an empirical distribution for $\hat{\beta}$ and $\hat{s^2}$ for each set of parameter values. From this, we calculate the mean and 95% confidence intervals of our distribution. These are displayed in Fig. S21. In each case, we set $\sigma = \sigma_1 = \cdots = \sigma_M$. Notice that the mean of the estimators for $\beta$ and $s^2$ are always in excellent agreement with the values (of $\beta$ and $s^2$, respectively) put in. Notice also that, although $s^2$ can never be negative, it is possible for the estimator $\hat{s^2}$ to be negative when there is a large amount of uncertainty in the estimation procedure (e.g. when $\sigma^2$ is large, Fig. S21h).

In the Supporting Information, we attach an `R` file, `hma_find_ssq_cis.R`, which enables the user to calculate an empirical mean and confidence intervals for $\hat{\beta}$ and $\hat{s}^2$ by using the sampling procedure just described. The user simply has to input values for $\beta$, $s^2$, $M$, and $\sigma_1, \ldots, \sigma_M$.

### Supplemental Tables

**Table S1.** Summary of contributed datasets.

| Species | Common Name | $M_{total}$ | $N_{total}$ | $\bar{N}$ | $s_N$ |
|---|---|---|---|---|---|
| *Alces alces* | Moose | 85 | 511951 | 6023 | 3229 |
| *Canis lupus* | Wolf | 29 | 145768 | 5026 | 3192 |
| *Cervus canadensis* | Elk | 25 | 97810 | 3912 | 1963 |
| *Didelphis virginiana* | Opossum | 30 | 43632 | 1454 | 985 |
| *Odocoileus hemionus* | Mule deer | 106 | 2455918 | 23169 | 8488 |
| *Odocoileus virginianus* | White-tailed deer | 28 | 610638 | 21809 | 4762 |
| *Ovibos moschatus* | Muskox | 36 | 782342 | 21732 | 14804 |
| *Rangifer tarandus caribou* | Caribou | 107 | 806482 | 7537 | 3504 |
| *Sus scrofa* | Pig | 35 | 163080 | 4659 | 3099 |
| *Tayassu pecari* | White-lipped peccary | 30 | 61002 | 2033 | 1448 |
| | *Totals* | 511 | 5678623 | 9736 | 4548 |

Note: $M_{total}$ = total number of animals in dataset; $N_{total}$ = total number of relocations in dataset; $\bar{N}$ = average number of relocations per animal; $s_N$ = standard deviation of relocations per animal. Totals are column sums for $M_{total}$ and $N_{total}$ and column means for $\bar{N}$ and $s_N$. All values under $\bar{N}$ and $s_N$ are rounded to the nearest whole number.

**Table S2.** RSF model structures for each species and landscape data information.

| Species | Model Structure (logit($p$) = ...) |
|---|---|
| *Alces alces* | $\text{Water}_c + \text{Fen}_c + \text{Sparse Treed}_c + \text{Deciduous}_c$ |
| *Canis lupus* | $\text{Water}_c + \text{Mixedwood}_c + \text{Jack Pine}_c + \text{Black Spruce}_c$ |
| *Cervus canadensis* | $\%\text{Mixedwood}_s + \text{Ruggedness}_s + \Delta\text{NDVI}_s + \text{Distance to Road}_s$ |
| *Didelphis virginiana* | $\text{Swamp}_c + \text{Evergreen}_c + \text{Deciduous}_c$ |
| *Odocoileus hemionus* | $\text{Temperature}_s + \text{Precipitation}_s + \text{Deciduous}_c + \text{Evergreen}_c + \text{Mixedwood}_c + \text{Shrub}_c$ |
| *Odocoileus virginianus* | $\text{Soybeans}_c + \text{Deciduous}_c + \text{Wetlands}_c$ |
| *Ovibos moschatus* | $\text{Snow/Ice}_c + \text{Sparse Vegetation}_c + \text{Dense Vegetation}_c + \text{Elevation}_s$ |
| *Rangifer tarandus caribou* | $\text{Forage}_s + \text{Mixedwood}_c + \text{Conifer}_c + \text{Disturbance}_c$ |
| *Sus scrofa* | $\text{Bottom}_c + \text{Evergreen}_c + \text{Crops}_c$ |
| *Tayassu pecari* | $\text{Canopy Closure}_s + \text{Distance to Road}_s + \text{Forest}_c + \text{Pasture}_c$ |

Note: A "c" subscript indicates the landscape predictor was a binary (categorical) raster layer. A "s" subscript indicates the landscape predictor was a centered-and-scaled (numeric) raster layer.

### Supplemental Figures

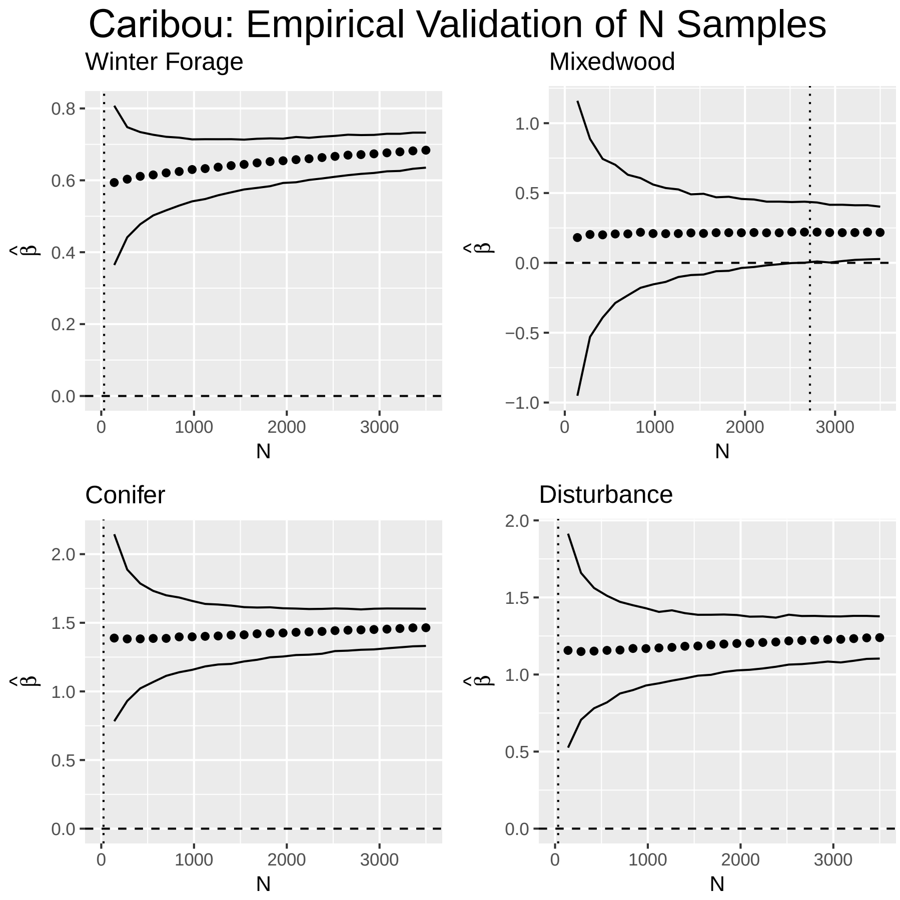


**Fig. S1.** Estimated selection coefficients and their empirical intervals with respect to $N$. Predicted $N$ ($N_{pred}$; vertical dotted line) strongly aligns with the value of $N$ at which the empirical interval of the selection coefficient ($\hat{\beta}$) no longer includes 0 ($N_{obs}$). If no vertical line occurs, then $N_{pred}$ exceeds the range of $N$.

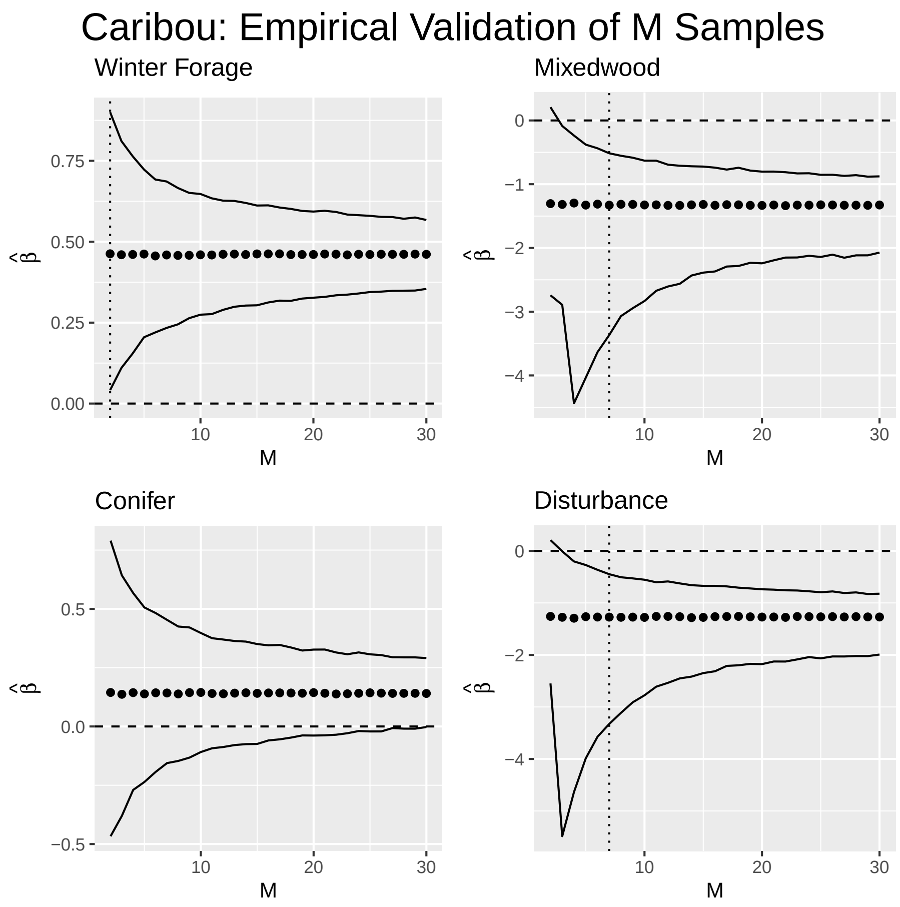


**Fig. S2.** Estimated selection coefficients and their empirical intervals with respect to $M$. Predicted $M$ ($M_{pred}$; vertical dotted line) strongly aligns with the value of $M$ at which the empirical interval of the selection coefficient ($\hat{\beta}$) no longer includes 0 ($M_{obs}$). If no vertical line occurs, then $M_{pred}$ exceeds the range of $M$.

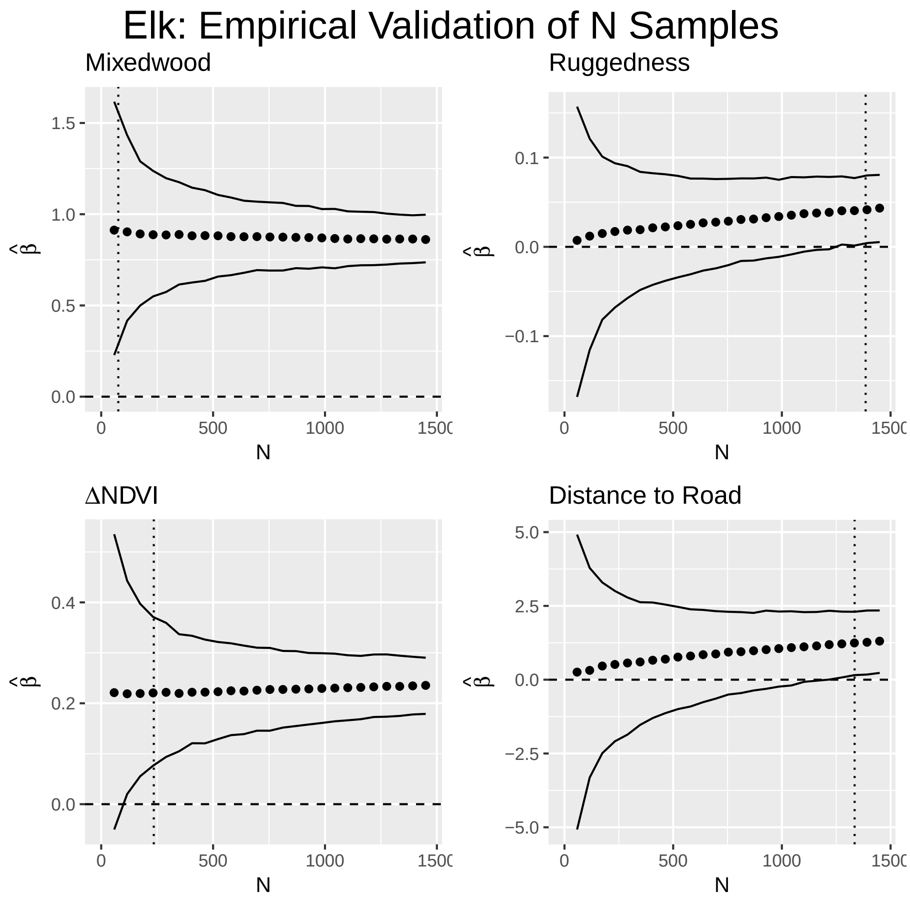


**Fig. S3.** Estimated selection coefficients and their empirical intervals with respect to $N$. Predicted $N$ ($N_{pred}$; vertical dotted line) strongly aligns with the value of $N$ at which the empirical interval of the selection coefficient ($\hat{\beta}$) no longer includes 0 ($N_{obs}$). If no vertical line occurs, then $N_{pred}$ exceeds the range of $N$.

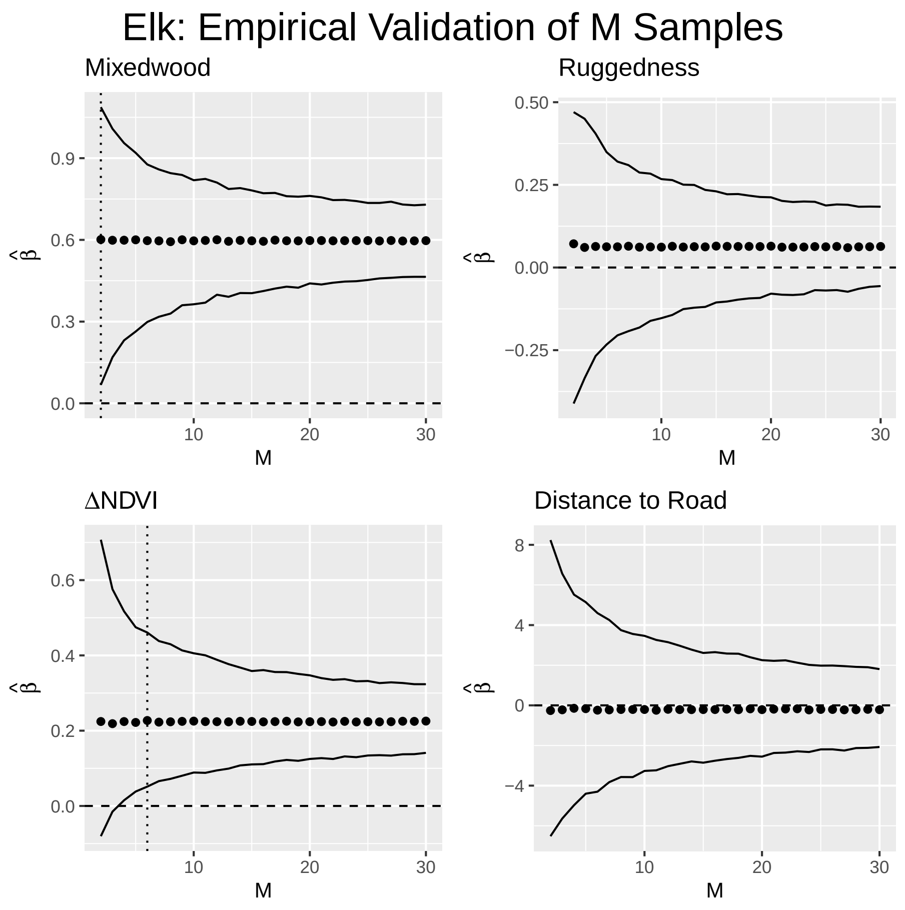


**Fig. S4.** Estimated selection coefficients and their empirical intervals with respect to $M$. Predicted $M$ ($M_{pred}$; vertical dotted line) strongly aligns with the value of $M$ at which the empirical interval of the selection coefficient ($\hat{\beta}$) no longer includes 0 ($M_{obs}$). If no vertical line occurs, then $M_{pred}$ exceeds the range of $M$.

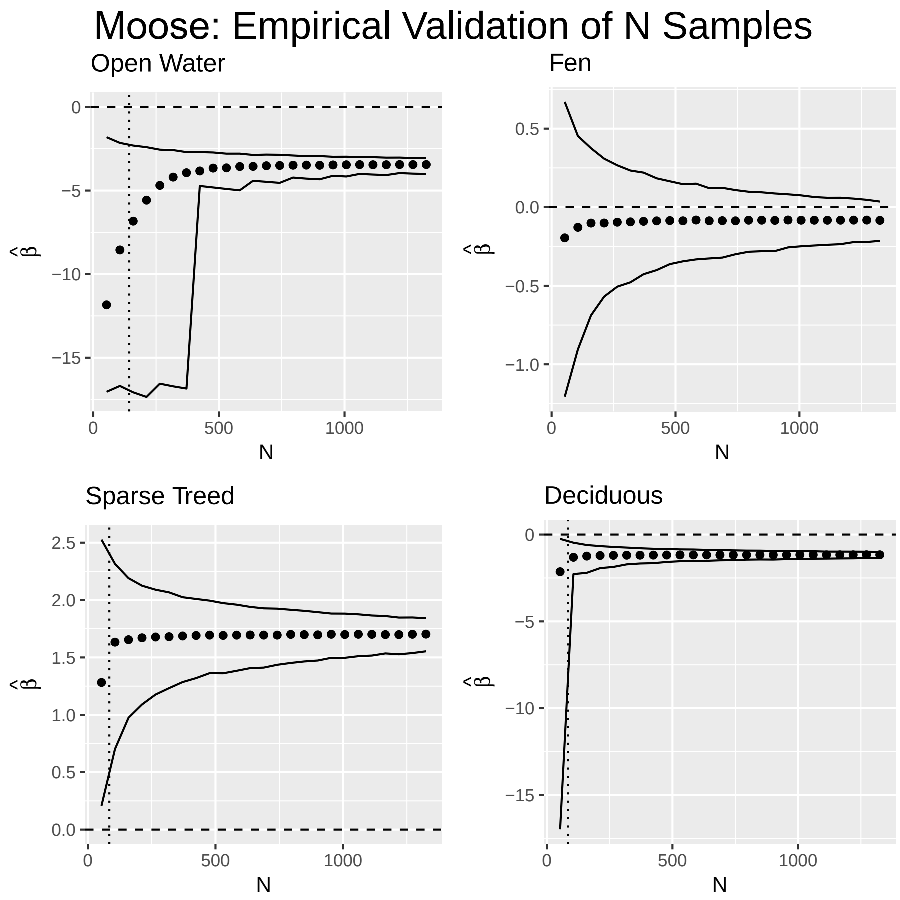


**Fig. S5.** Estimated selection coefficients and their empirical intervals with respect to $N$. Predicted $N$ ($N_{pred}$; vertical dotted line) strongly aligns with the value of $N$ at which the empirical interval of the selection coefficient ($\hat{\beta}$) no longer includes 0 ($N_{obs}$). If no vertical line occurs, then $N_{pred}$ exceeds the range of $N$.

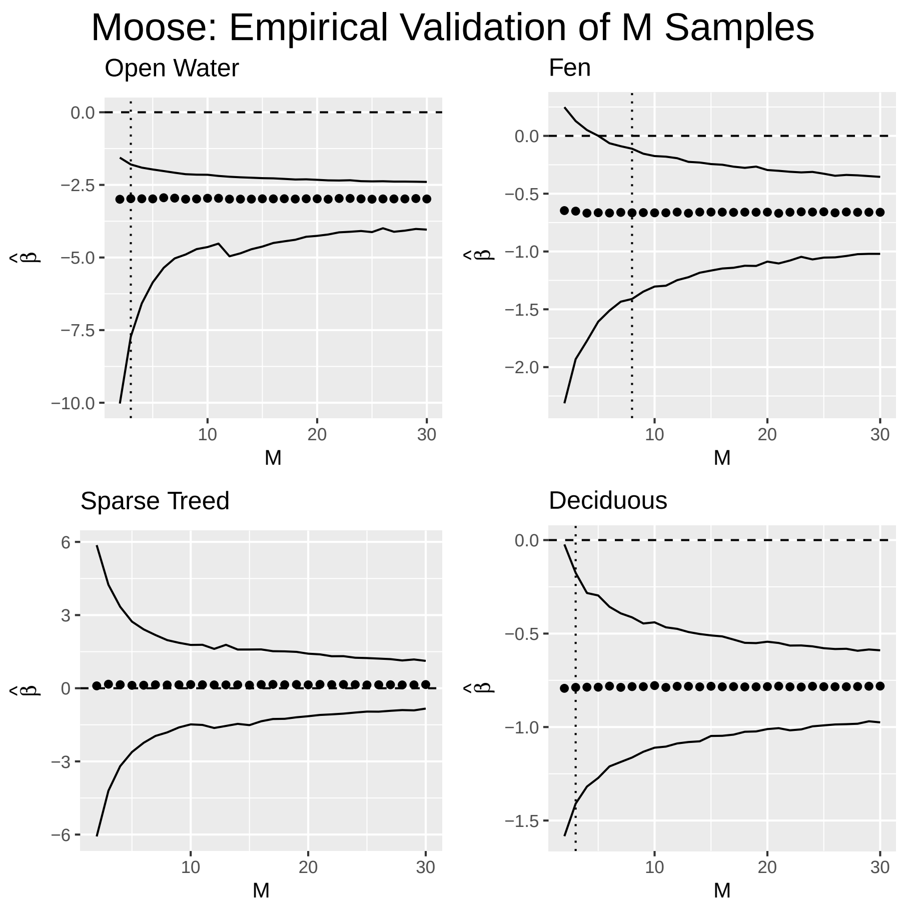


**Fig. S6.** Estimated selection coefficients and their empirical intervals with respect to $M$. Predicted $M$ ($M_{pred}$; vertical dotted line) strongly aligns with the value of $M$ at which the empirical interval of the selection coefficient ($\hat{\beta}$) no longer includes 0 ($M_{obs}$). If no vertical line occurs, then $M_{pred}$ exceeds the range of $M$.

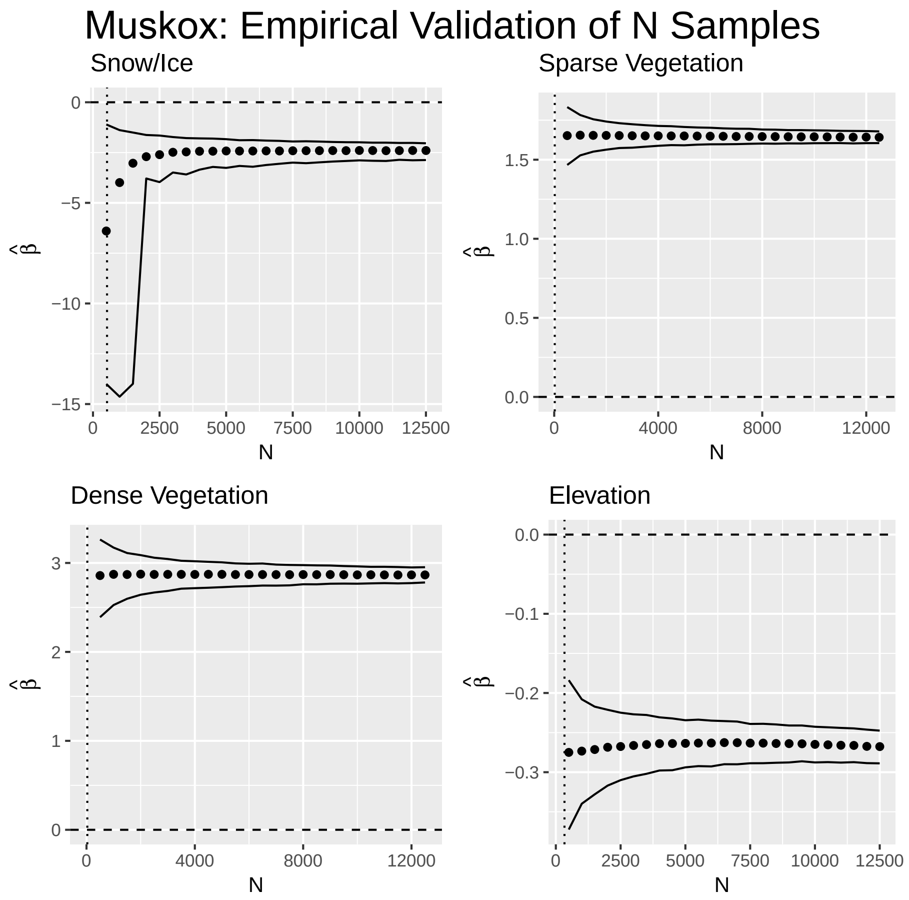


**Fig. S7.** Estimated selection coefficients and their empirical intervals with respect to $N$. Predicted $N$ ($N_{pred}$; vertical dotted line) strongly aligns with the value of $N$ at which the empirical interval of the selection coefficient ($\hat{\beta}$) no longer includes 0 ($N_{obs}$). If no vertical line occurs, then $N_{pred}$ exceeds the range of $N$.

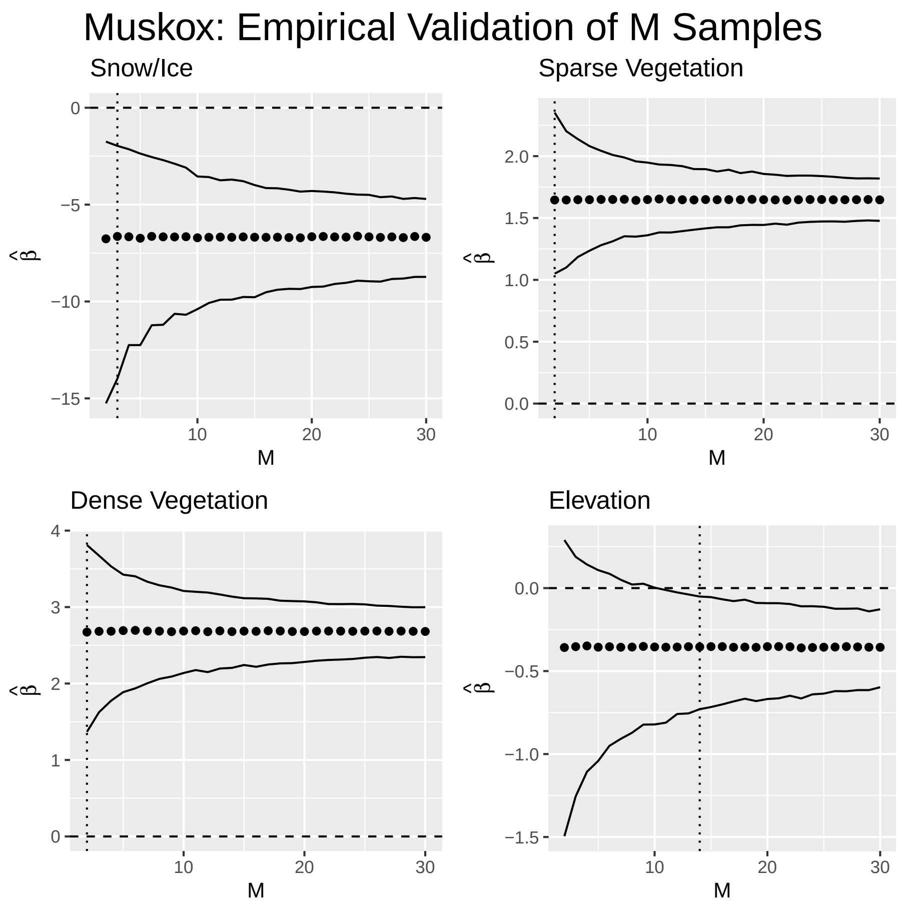


**Fig. S8.** Estimated selection coefficients and their empirical intervals with respect to $M$. Predicted $M$ ($M_{pred}$; vertical dotted line) strongly aligns with the value of $M$ at which the empirical interval of the selection coefficient ($\hat{\beta}$) no longer includes 0 ($M_{obs}$). If no vertical line occurs, then $M_{pred}$ exceeds the range of $M$.

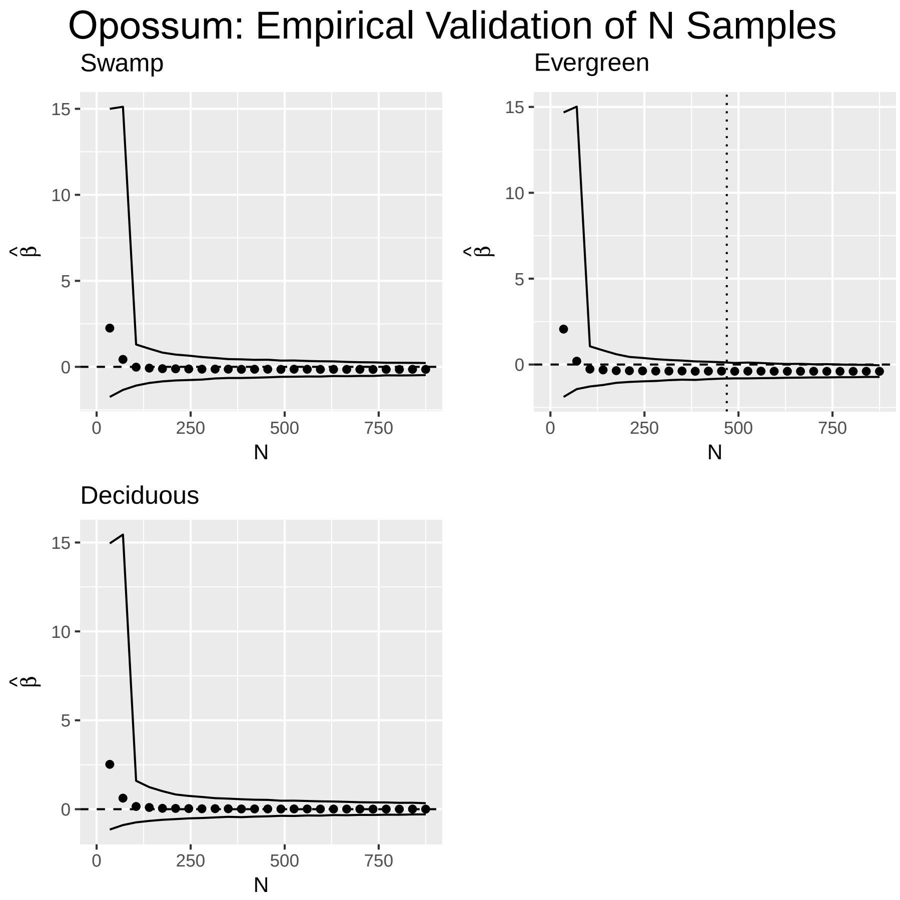


**Fig. S9.** Estimated selection coefficients and their empirical intervals with respect to $N$. Predicted $N$ ($N_{pred}$; vertical dotted line) strongly aligns with the value of $N$ at which the empirical interval of the selection coefficient ($\hat{\beta}$) no longer includes 0 ($N_{obs}$). If no vertical line occurs, then $N_{pred}$ exceeds the range of $N$.

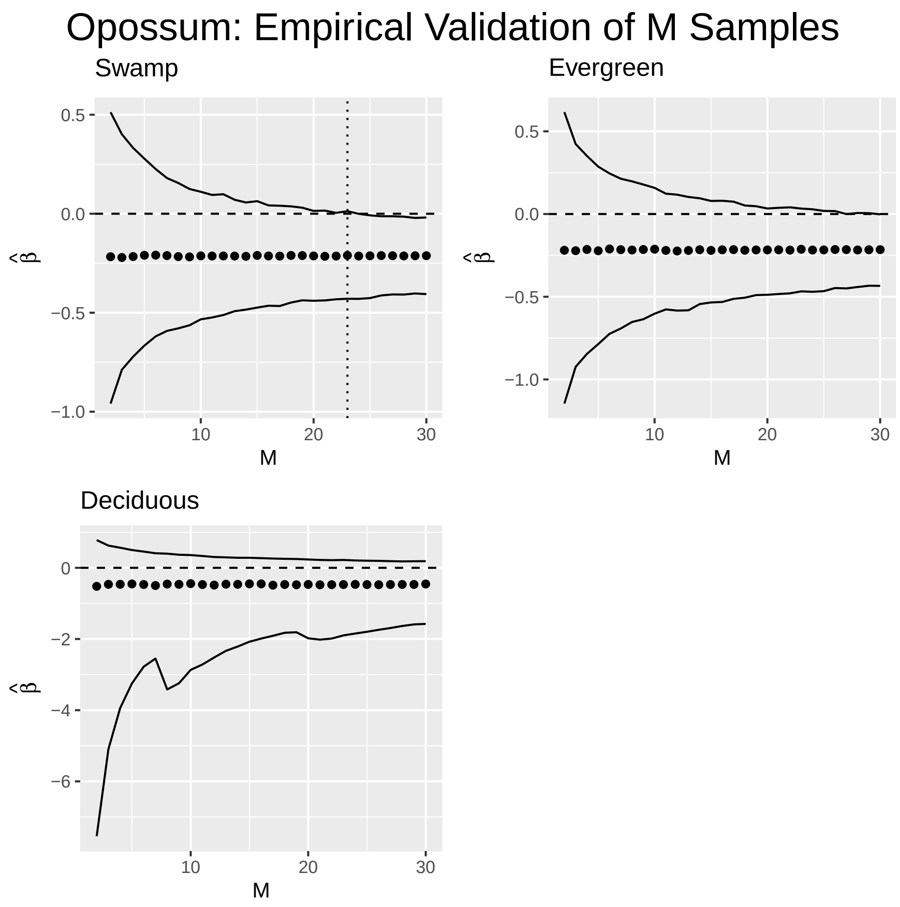


**Fig. S10.** Estimated selection coefficients and their empirical intervals with respect to $M$. Predicted $M$ ($M_{pred}$; vertical dotted line) strongly aligns with the value of $M$ at which the empirical interval of the selection coefficient ($\hat{\beta}$) no longer includes 0 ($M_{obs}$). If no vertical line occurs, then $M_{pred}$ exceeds the range of $M$.

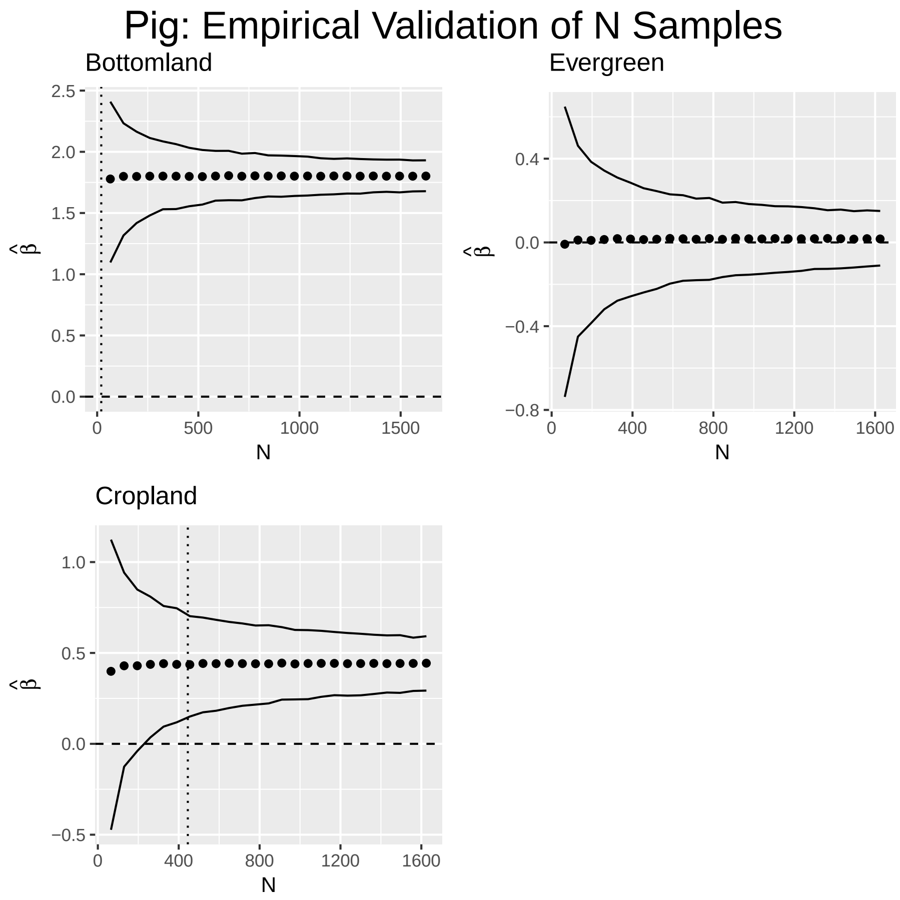


**Fig. S11.** Estimated selection coefficients and their empirical intervals with respect to $N$. Predicted $N$ ($N_{pred}$; vertical dotted line) strongly aligns with the value of $N$ at which the empirical interval of the selection coefficient ($\hat{\beta}$) no longer includes 0 ($N_{obs}$). If no vertical line occurs, then $N_{pred}$ exceeds the range of $N$.

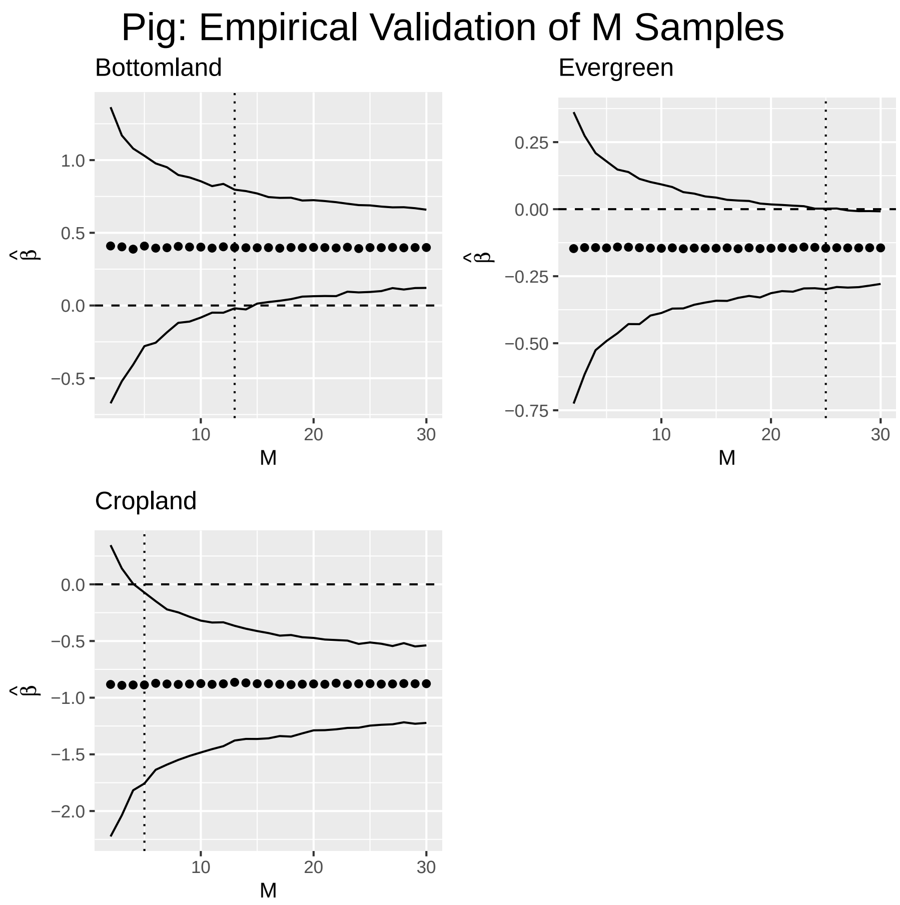


**Fig. S12.** Estimated selection coefficients and their empirical intervals with respect to $M$. Predicted $M$ ($M_{pred}$; vertical dotted line) strongly aligns with the value of $M$ at which the empirical interval of the selection coefficient ($\hat{\beta}$) no longer includes 0 ($M_{obs}$). If no vertical line occurs, then $M_{pred}$ exceeds the range of $M$.

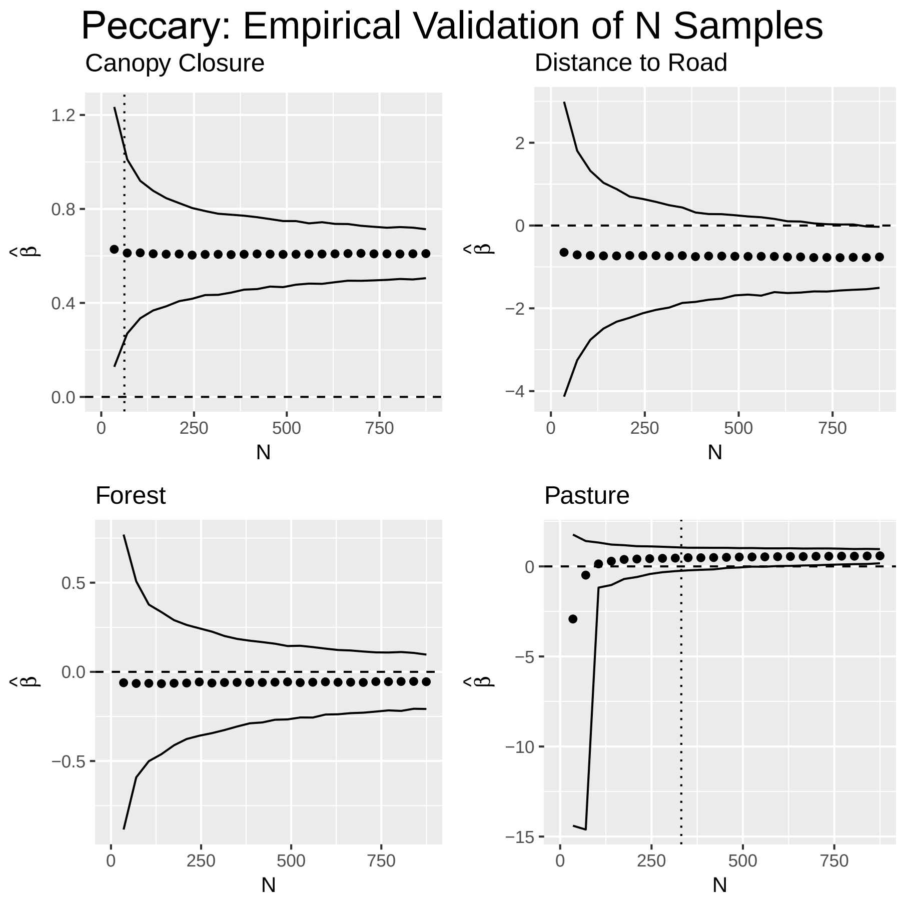


**Fig. S13.** Estimated selection coefficients and their empirical intervals with respect to $N$. Predicted $N$ ($N_{pred}$; vertical dotted line) strongly aligns with the value of $N$ at which the empirical interval of the selection coefficient ($\hat{\beta}$) no longer includes 0 ($N_{obs}$). If no vertical line occurs, then $N_{pred}$ exceeds the range of $N$.

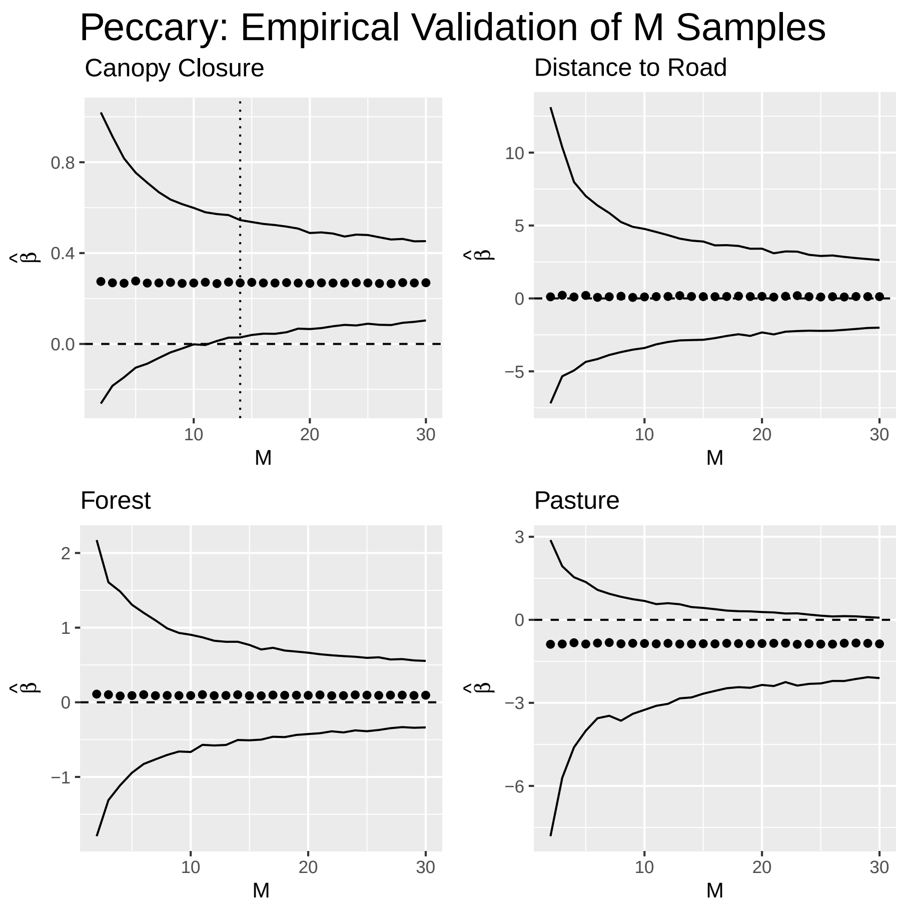


**Fig. S14.** Estimated selection coefficients and their empirical intervals with respect to $M$. Predicted $M$ ($M_{pred}$; vertical dotted line) strongly aligns with the value of $M$ at which the empirical interval of the selection coefficient ($\hat{\beta}$) no longer includes 0 ($M_{obs}$). If no vertical line occurs, then $M_{pred}$ exceeds the range of $M$.

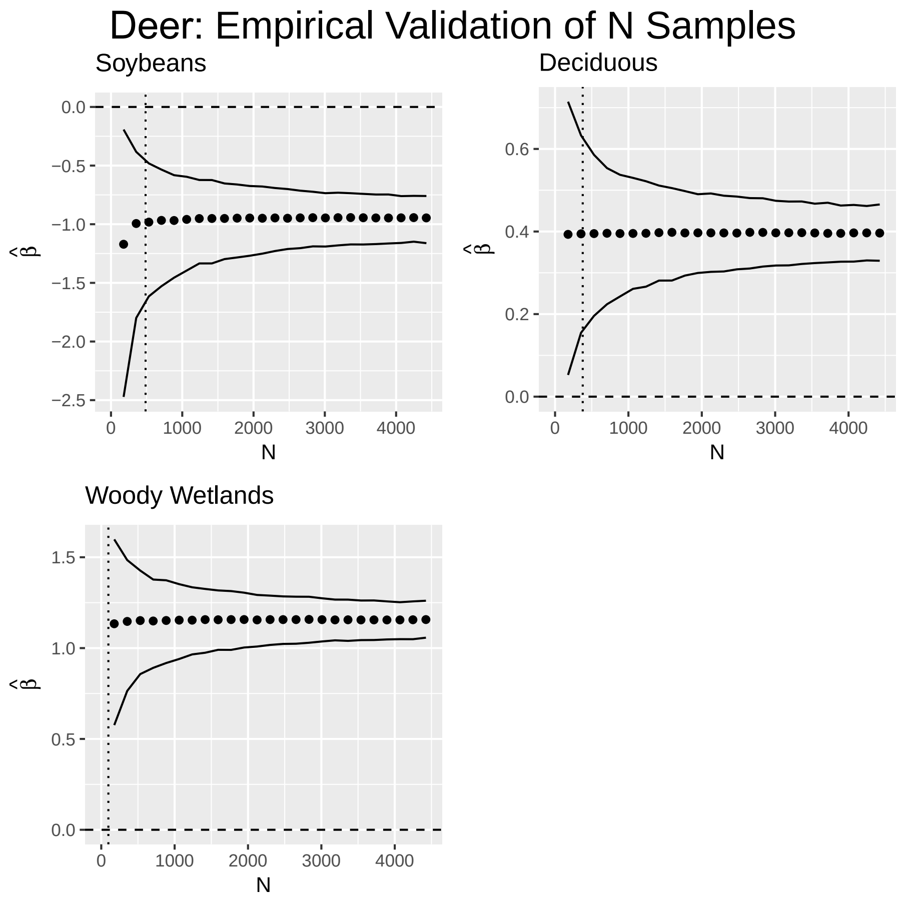


**Fig. S15.** Estimated selection coefficients and their empirical intervals with respect to $N$. Predicted $N$ ($N_{pred}$; vertical dotted line) strongly aligns with the value of $N$ at which the empirical interval of the selection coefficient ($\hat{\beta}$) no longer includes 0 ($N_{obs}$). If no vertical line occurs, then $N_{pred}$ exceeds the range of $N$.

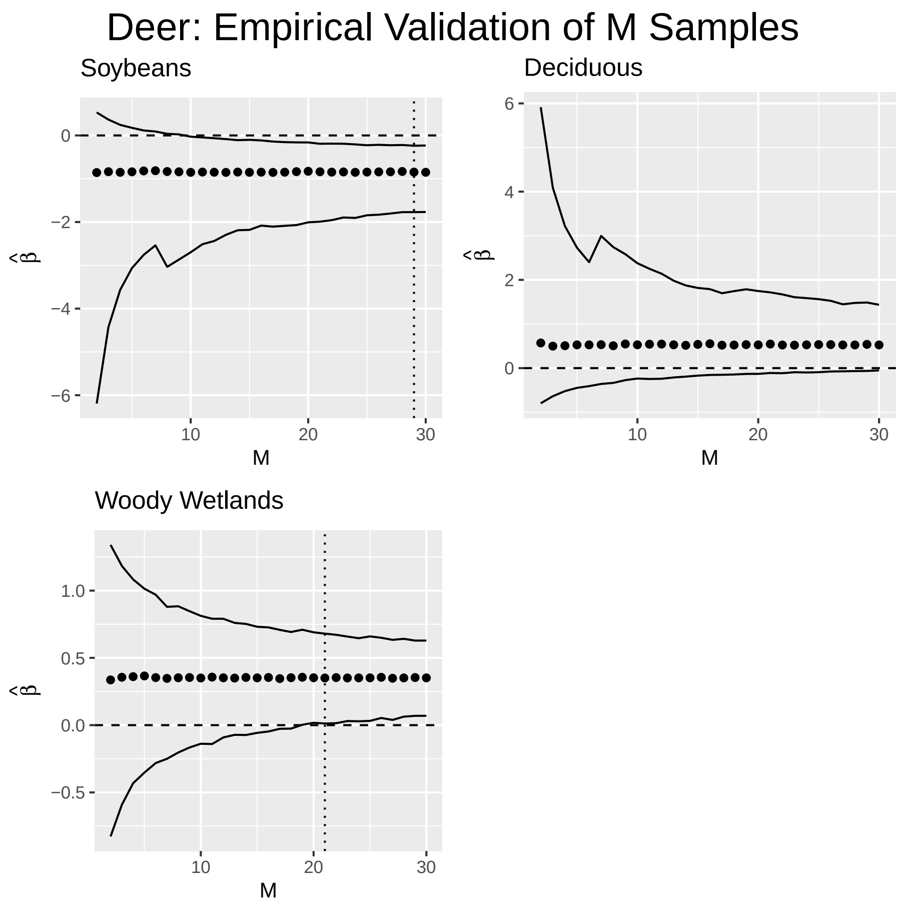


**Fig. S16.** Estimated selection coefficients and their empirical intervals with respect to $M$. Predicted $M$ ($M_{pred}$; vertical dotted line) strongly aligns with the value of $M$ at which the empirical interval of the selection coefficient ($\hat{\beta}$) no longer includes 0 ($M_{obs}$). If no vertical line occurs, then $M_{pred}$ exceeds the range of $M$.

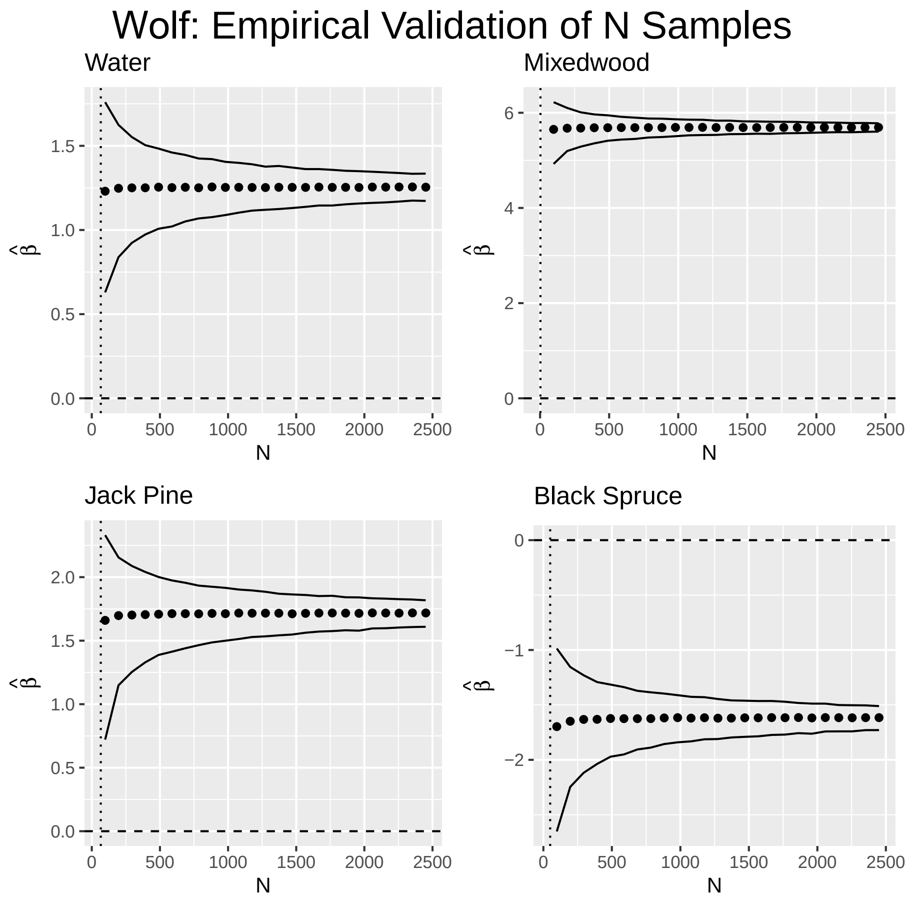


**Fig. S17.** Estimated selection coefficients and their empirical intervals with respect to $N$. Predicted $N$ ($N_{pred}$; vertical dotted line) strongly aligns with the value of $N$ at which the empirical interval of the selection coefficient ($\hat{\beta}$) no longer includes 0 ($N_{obs}$). If no vertical line occurs, then $N_{pred}$ exceeds the range of $N$.

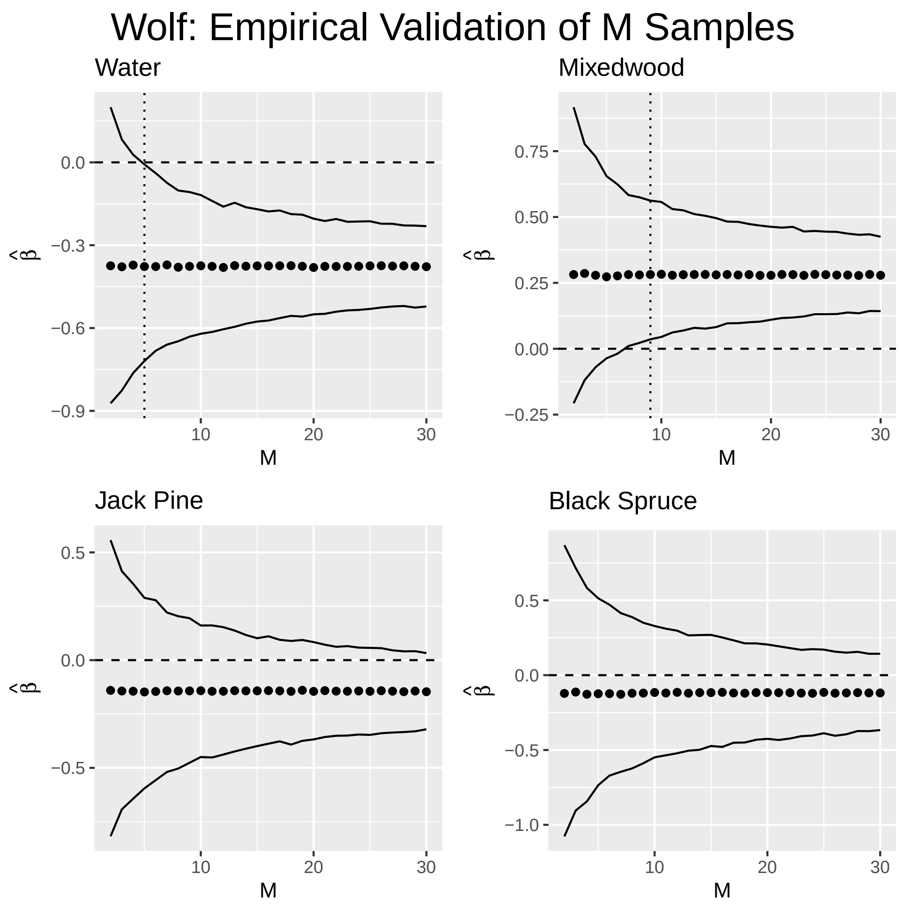


**Fig. S18.** Estimated selection coefficients and their empirical intervals with respect to $M$. Predicted $M$ ($M_{pred}$; vertical dotted line) strongly aligns with the value of $M$ at which the empirical interval of the selection coefficient ($\hat{\beta}$) no longer includes 0 ($M_{obs}$). If no vertical line occurs, then $M_{pred}$ exceeds the range of $M$.

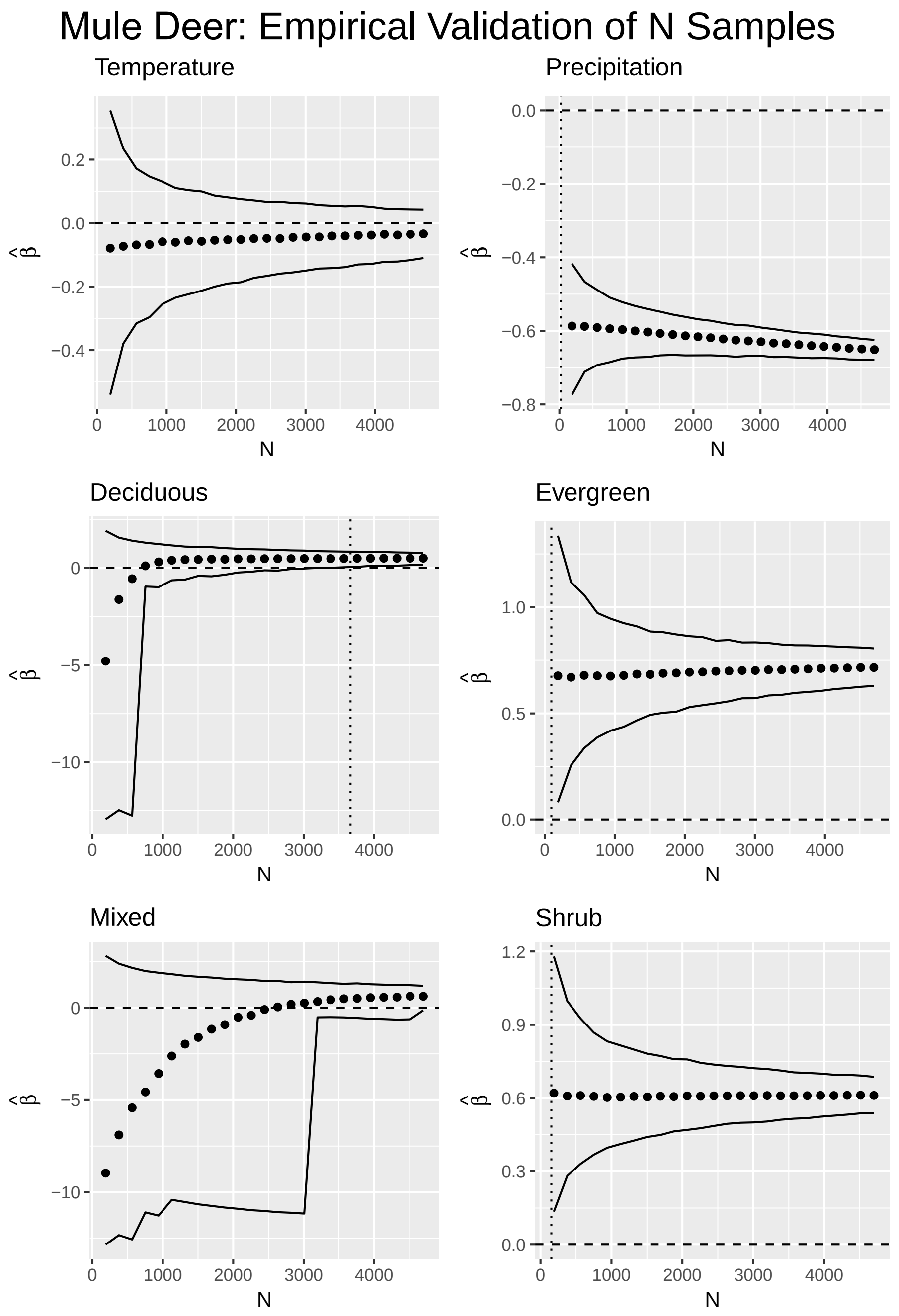


**Fig. S19.** Estimated selection coefficients and their empirical intervals with respect to $N$. Predicted $N$ ($N_{pred}$; vertical dotted line) strongly aligns with the value of $N$ at which the empirical interval of the selection coefficient ($\hat{\beta}$) no longer includes 0 ($N_{obs}$). If no vertical line occurs, then $N_{pred}$ exceeds the range of $N$.

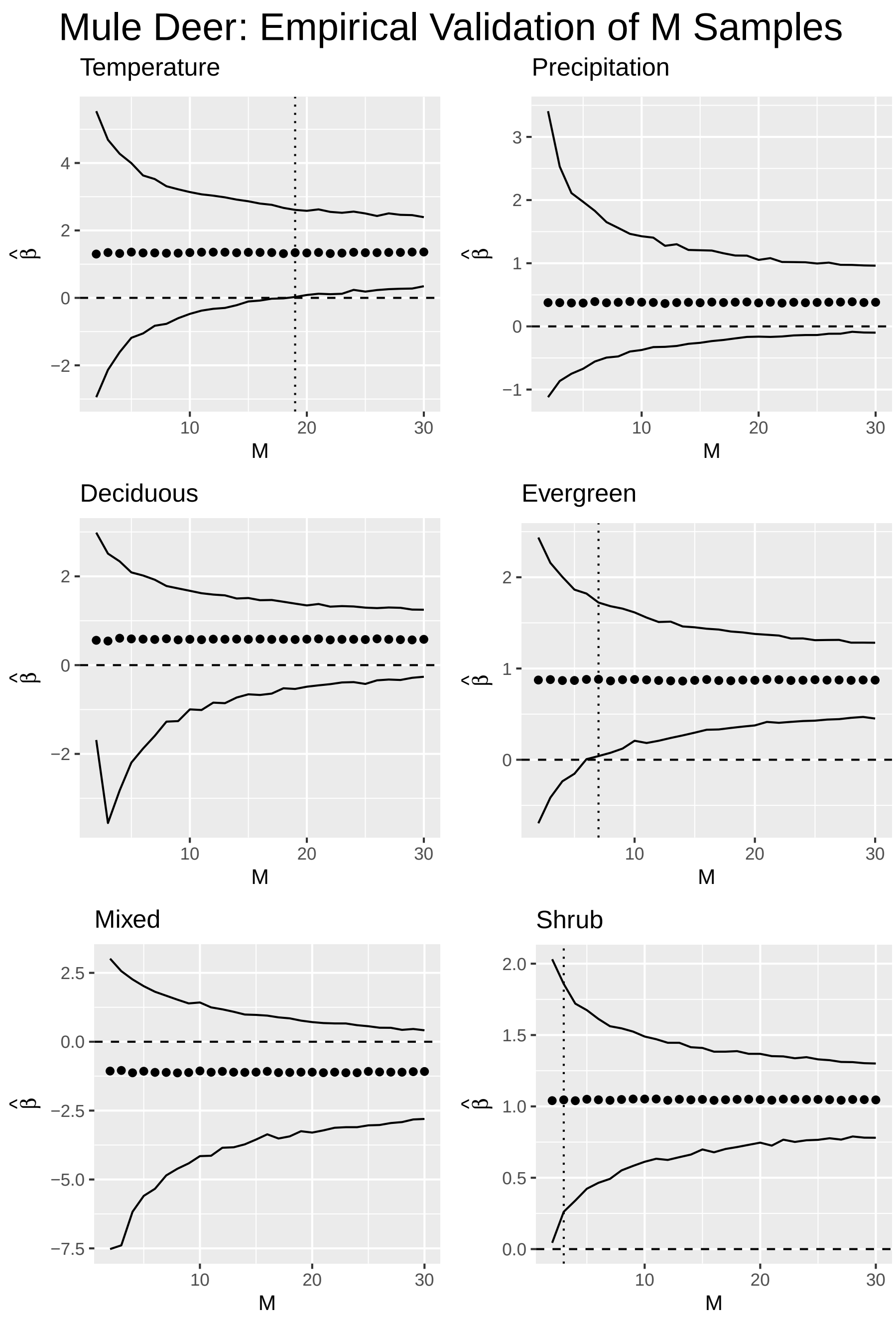


**Fig. S20.** Estimated selection coefficients and their empirical intervals with respect to $M$. Predicted $M$ ($M_{pred}$; vertical dotted line) strongly aligns with the value of $M$ at which the empirical interval of the selection coefficient ($\hat{\beta}$) no longer includes 0 ($M_{obs}$). If no vertical line occurs, then $M_{pred}$ exceeds the range of $M$.

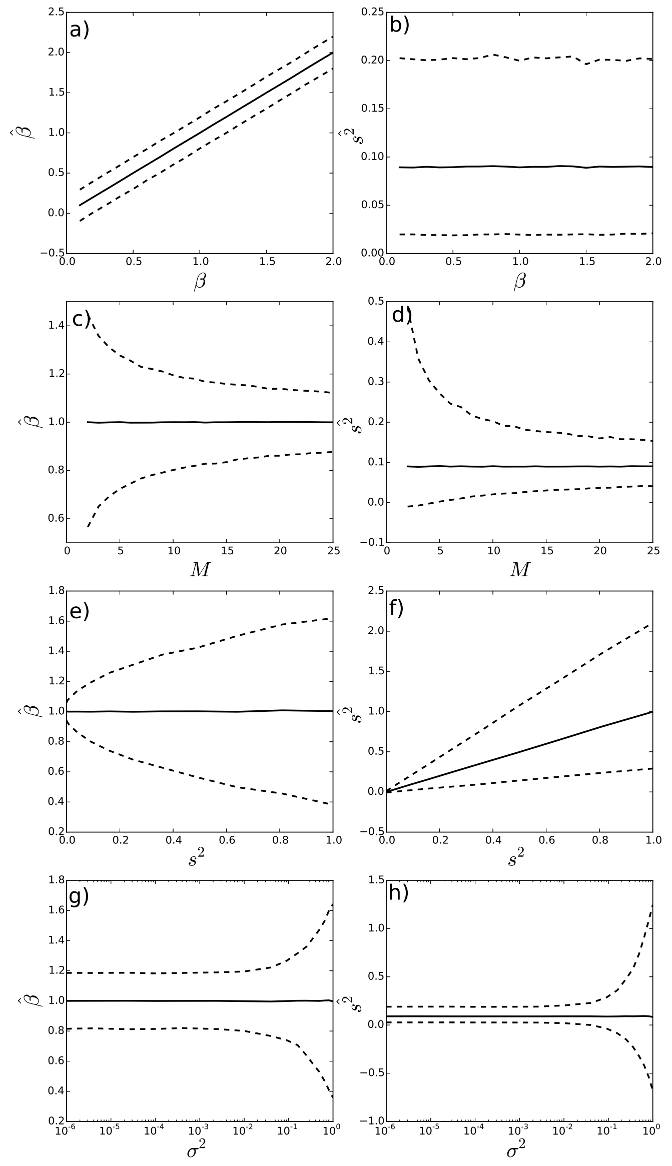


**Fig. S21. Mean and 95% confidence intervals of estimators for $\beta$ and $s^2$.** Each panel in the left-hand column shows the empirical mean (solid curves) and 95% confidence intervals (dashed curves) of $\hat{\beta}$ for different values of $\beta$, $s^2$, $M$, and $\sigma^2$. The panels in right-hand column show the same for $\hat{s^2}$. Unless otherwise stated, $\beta = 1$, $s^2 = 0.09$, $\sigma^2 = 0.01$, $M = 10$.
