## Supplementary material for "Solving the Sample Size Problem for Resource Selection Analysis": Finding Sums of Squares R Supplement

rm(list=ls());gc() # Clear the memory

##########################################################################
# Input values - beta, s, M, sigma, alpha
#
# Edit these to the values you require
#  - beta: The estimated value of beta, the mean selection strength across 
#          animals in the population
#  - s_val: The estimated value of s, the standard deviation of the selection
#           strength across all animals in the population
#  - M_val: Number of animals tagged
#  - sigmas: A list [sigma_1,sigma_2,...,sigma_M] of the sigma_i values for
#            each individual (this can be derived from the equation for
#            sigma_i in the text
#  - alpha: the desired confidence level.  The output will report the 
#           alpha/2 and (1-alpha/2) quantiles
##########################################################################
beta<-1
s_val<-0.3
M_val<-10
sigmas<-rep(0.1,M_val)
alpha<-0.05

##########################################################################
# Number of samples should be so that there are at least 100 samples in 
# each tail
##########################################################################
no_samples<-ceiling(200/alpha)

##########################################################################
# Set up empty arrays for hat_beta and hat_ssq (estimators of beta and 
# s^2, respectively)
##########################################################################
hat_beta<-c()
hat_ssq<-c()

for (sample in 1:no_samples)
{
  ##########################################################################
  # Sample beta_i from N(beta,s^2) for each i
  ##########################################################################
  betais<-rnorm(M_val,beta,s_val)
  
  ##########################################################################
  # Sample hat_beta_i from N(beta_i, sigma^2) for each i
  ##########################################################################
  hat_betais<-rnorm(M_val,betais,sigmas)

  ##########################################################################
  # Compute hat_beta and add to list
  ##########################################################################
  hat_beta<-c(hat_beta,mean(hat_betais))

  ##########################################################################
  # Compute hat_ssq and add to list
  ##########################################################################
  hat_ssq<-c(hat_ssq,var(hat_betais)-mean(sigmas^2))
}

##########################################################################
# Get median, mean, SD and 95% confidence intervals for hat_beta and hat_ssq
##########################################################################
mean_beta<-mean(hat_beta)
sorted_hat_beta<-sort(hat_beta)
lower_beta<-sorted_hat_beta[no_samples*alpha/2]
upper_beta<-sorted_hat_beta[no_samples*(1-alpha/2)]

mean_ssq<-mean(hat_ssq)
sorted_hat_ssq<-sort(hat_ssq)
lower_ssq<-sorted_hat_ssq[no_samples*alpha/2]
upper_ssq<-sorted_hat_ssq[no_samples*(1-alpha/2)]

##########################################################################
# Print mean, lower and upper quantiles for beta and s^2
##########################################################################
mean_beta
lower_beta
upper_beta
mean_ssq
lower_ssq
upper_ssq
